# Development and characterization of phospho-ubiquitin antibodies to monitor PINK1-PRKN signaling in cells and tissue

**DOI:** 10.1101/2024.01.15.575715

**Authors:** Jens O. Watzlawik, Xu Hou, Tyrique Richardson, Szymon L. Lewicki, Joanna Siuda, Zbigniew K. Wszolek, Casey N. Cook, Leonard Petrucelli, Michael DeTure, Dennis W. Dickson, Odetta Antico, Miratul M. K. Muqit, Jordan B. Fishman, Karima Pirani, Ravindran Kumaran, Nicole K. Polinski, Fabienne C. Fiesel, Wolfdieter Springer

**Affiliations:** Department of Neuroscience, Mayo Clinic, Jacksonville, FL 32224, USA; Department of Neurology, Faculty of Medical Sciences in Katowice, Medical University of Silesia, Katowice 40-055, Poland; Department of Neurology, Mayo Clinic, Jacksonville, FL 32224, USA; Neuroscience PhD Program, Mayo Clinic Graduate School of Biomedical Sciences, Jacksonville, FL 32224, USA; MRC Protein Phosphorylation and Ubiquitylation Unit, School of Life Sciences, University of Dundee, Dundee, DD1 5EH, United Kingdom; 21st Century Biochemicals Inc., Marlborough, MA 01752, USA; ImmunoPrecise Antibodies Ltd., Victoria, BC V8Z 7X8, Canada; Abcam plc, Cambridge, CB2 0AX, United Kingdom; The Michael J. Fox Foundation for Parkinson’s Research, New York, NY 10163, USA

**Author notes:** Correspondence should be addressed to: Wolfdieter Springer, PhD, Department of Neuroscience; Mayo Clinic, 4500 San Pablo Road, Jacksonville, FL 32224, USA. Contributed equally (Jens O. Watzlawik, Xu Hou).

**Keywords:** autophagy, mitochondria, mitophagy, Parkin, Parkinson disease, phospho-ubiquitin, PINK1, PRKN, recombinant antibody, ubiquitin

## Abstract

The selective removal of dysfunctional mitochondria, a process termed mitophagy, is critical for cellular health and impairments have been linked to aging, Parkinson disease, and other neurodegenerative conditions. A central mitophagy pathway is orchestrated by the ubiquitin (Ub) kinase PINK1 together with the E3 Ub ligase PRKN/Parkin. The decoration of damaged mitochondrial domains with phosphorylated Ub (p-S65-Ub) mediates their elimination though the autophagy system. As such p-S65-Ub has emerged as a highly specific and quantitative marker of mitochondrial damage with significant disease relevance. Existing p-S65-Ub antibodies have been successfully employed as research tools in a range of applications including western blot, immunocytochemistry, immunohistochemistry, and ELISA. However, physiological levels of p-S65-Ub in the absence of exogenous stress are very low, therefore difficult to detect and require reliable and ultrasensitive methods. Here we generated and characterized a collection of novel recombinant, rabbit monoclonal p-S65-Ub antibodies with high specificity and affinity in certain applications that allow the field to better understand the molecular mechanisms and disease relevance of PINK1-PRKN signaling. These antibodies may also serve as novel diagnostic or prognostic tools to monitor mitochondrial damage in various clinical and pathological specimens.

## INTRODUCTION

Mitophagy is a cytoprotective mechanism for the selective and timely removal of dysfunctional or superfluous mitochondria through autophagy. One central mitophagy mechanism that provides specificity for damaged mitochondria is the phospho-ubiquitylation of mitochondrial proteins, which fosters the recruitment of autophagy receptors only to those organelles that need to be degraded [1]. PTEN-induced kinase 1 (PINK1) and the E3 ubiquitin (Ub) ligase Parkin (PRKN) are the key players of this Ub-tagging process and complete loss of either enzyme leads to early-onset Parkinson disease (PD). Upon mitochondrial damage, PINK1 locally stabilizes and phosphorylates serine 65 on Ub (p-S65-Ub) that is either free or already attached to outer mitochondrial proteins as monomer or different types of poly-Ub chains [2,3]. PINK1 also phosphorylates the E3 Ub ligase PRKN at a conserved serine at position 65 within its Ub-like domain [4,5]. Both PRKN phosphorylation and allosteric p-S65-Ub binding fully activates its E3 Ub ligase activity [6–8]. Activated PRKN attaches additional Ub moieties to mitochondrial proteins, which can be further phosphorylated by PINK1 and serve as additional docking stations to recruit even more cytosolic PRKN to mitochondria. This positive feedback loop between PINK1 and PRKN amplifies the extent of p-S65-Ub coating of damaged mitochondria, which drives their elimination via mitophagy.

Given the causal link between the loss of function in PINK1 or PRKN and early-onset PD, mitophagy dysfunction has been long considered to play an important role in disease pathogenesis [9]. Mitochondrial impairments are commonly found in aging and age-related diseases including neurodegenerative disorders, and mitophagy defects are likely also widespread [10]. As the joint product of PINK1 and PRKN enzymatic activity, p-S65-Ub can be used as a quantitative measure of mitochondrial health and mitophagy alteration. Absent or reduced p-S65-Ub levels result from failure to properly activate mitophagy in model organisms with *PINK1* or *PRKN* knockout or mutations [11–13]. In contrast, elevated p-S65-Ub levels can be detected with age and in many diseases and may originate from increased mitochondrial damage or rather may accumulate due to a downstream block of the autophagic-lysosomal degradation system [11,13–15]. p-S65-Ub therefore represents a promising and potent diagnostic marker for damaged mitochondria in cell and animal models as well as in human biofluids and postmortem brain [11,13]. Indeed, p-S65-Ub is currently being tested in several preclinical studies as an early biomarker in different neurodegenerative diseases. Existing p-S65-Ub antibodies show encouraging results in ELISA-based applications. However, the overall p-S65-Ub signal in human biofluids at baseline and under pathological conditions is very low and therefore difficult to detect in a reliable manner. Similarly, sensitive histopathological assessment of damaged mitochondria undergoing mitophagy in human brain is currently very limited and would greatly benefit from more sensitive and reliable tools to further validate injury-prone brain regions and subcellular locations that are positive for the p-S65-Ub signal. In general, the detection of endogenous p-S65-Ub *in vivo* at baseline or under low (chronic) stress is key and requires more sensitive techniques and antibodies.

To expand the existing repertoire of tools, we here generated and thoroughly characterized a large series of recombinant, rabbit monoclonal p-S65-Ub antibodies (**Fig. 1**). The top performing p-S65-Ub antibody clones for each category were affinity-purified and further interrogated in cells and brain tissue of mouse and human origin and across a range of different applications. The top antibody candidates target p-S65-Ub with high specificity and affinity in different applications and are available as a novel resource for the community to study PINK1-PRKN signaling under normal, stress, or disease conditions.

**Figure 1.**
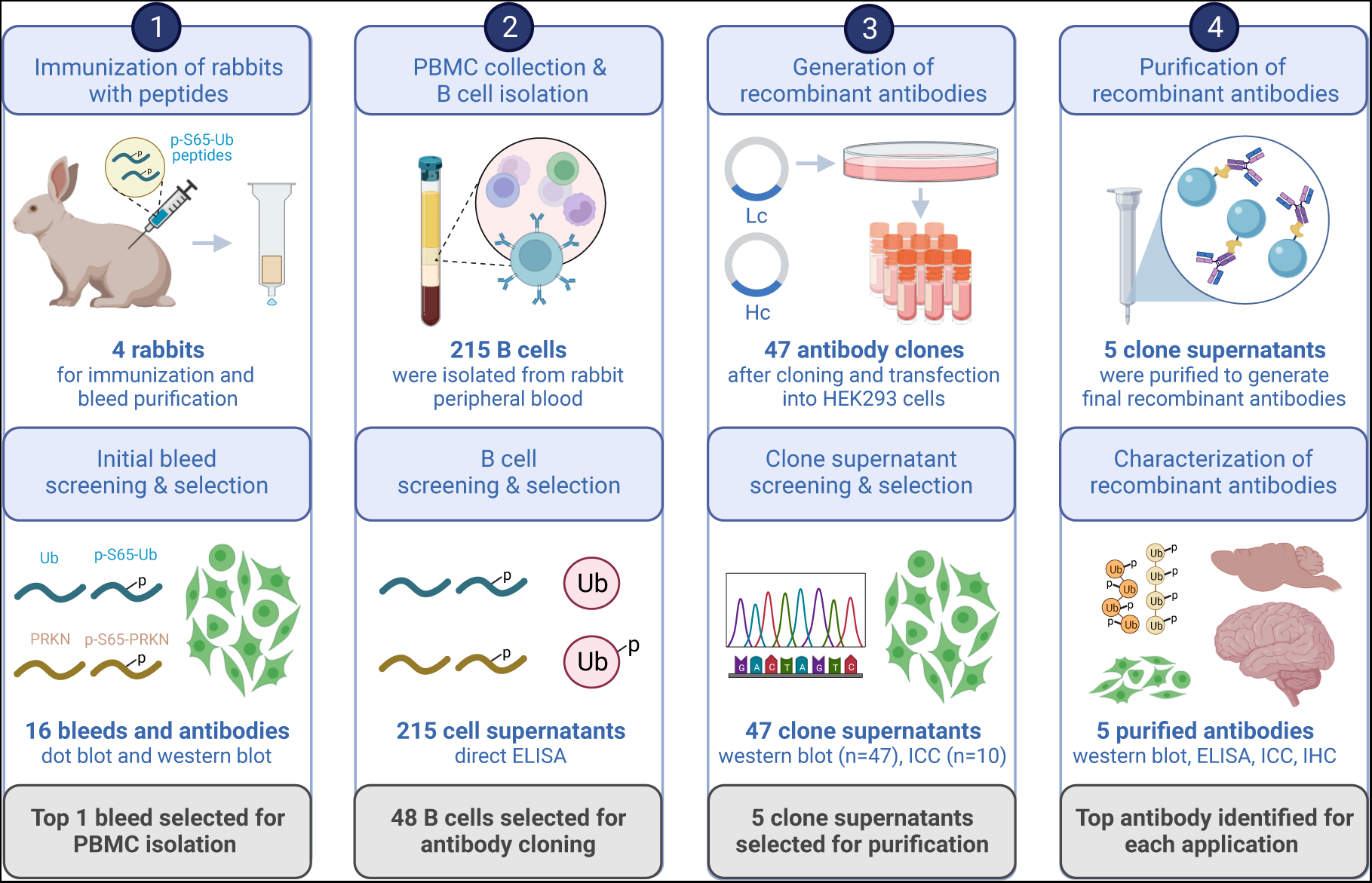
Schematic overview of recombinant p-S65-Ub antibody clone generation, selection, and validation. The illustration summarizes the entire process of recombinant, rabbit monoclonal antibody production starting from testing of the initial polyclonal bleeds to the isolation and screening of B cells, and the subsequent cloning of variable heavy and light chain antibody regions into an IgG vector frame. Recombinant antibodies were then sequenced, expressed in HEK293 cells and supernatants were screened by western blot and immunocytochemistry. The five most promising antibody clones were affinity purified and re-validated by a series of biochemical and imaging analyses using recombinant proteins as well as cells and brain tissues from mouse and human origin. PBMC - peripheral blood mononuclear cell, Lc - light chain, Hc - heavy chain, ICC - immunocytochemistry, IHC – immunohistochemistry. Created with BioRender.com.

## RESULTS

### Generation and screening of polyclonal rabbit antibodies

Four rabbits were immunized with a combination of N- or C-terminally conjugated p-S65-Ub peptides. Initial bleeds from these animals were tested for immunoreactivity in dot blots towards the respective phosphorylated- and non-phosphorylated Ub and PRKN peptides (**Fig. S1A**). Based on their stronger immunoreactivity, rabbits (#15021 and #15022) were further selected to receive additional p-S65-Ub peptide booster injections. The resulting affinity-purified polyclonal antibodies were tested in western blots using wild-type (WT) and negative control *PINK1* knockout (KO) human embryonic kidney (HEK293E) cells that were each treated with the mitochondrial stressor carbonyl cyanide 3-chlorophenylhydrazone (CCCP) **(Fig. S1B)** [13]. Affinity purified bleeds from rabbit #15022 but not rabbit #15021 demonstrated strong p-S65-Ub immunoreactivity in CCCP-treated WT cells, which was comparable to the commercially available reference antibody. Additional immuno-depletion further reduced the non-specific signal in *PINK1* KO cells. Given the superior signal-to-noise ratio, rabbit #15022 was selected for isolation of peripheral blood mononuclear cells (PBMCs).

### Recombinant monoclonal antibody generation and clonal selection

Collected PBMCs were enriched for antigen-specific B cells, and subsequently diluted for single B cell cultivation. Cell supernatants from a total of 215 single B cell cultures were tested by direct ELISA for immunoreactivity towards phosphorylated (serine 65) and non-phosphorylated full-length, monomeric Ub protein as well as non-/phosphorylated Ub and PRKN peptides surrounding the respective serine 65 residues (**Table S1**). We identified 48 candidates that consisted of 36 B cell cultures producing antibodies that strongly targeted both p-S65-Ub protein and peptide, four with similar specificity but weaker immunoreactivity, four that detected p-S65-Ub protein only but not peptide, and another four that also recognized the corresponding p-S65-PRKN peptide (**Table S1**). Variable regions of both heavy and kappa chains were subsequently cloned, sequenced, and co-expressed in HEK293 cells to give rise to 47 recombinant IgG antibody clones. A phylogenetic tree was created to determine their clonal relationship (**Fig. S2**).

We next screened all 47 recombinant clonal supernatants in western blots for affinity and selectivity (**Fig. S3**). Ten supernatants showed substantially stronger signal with lysates from CCCP-treated WT HEK293E compared to *PINK1* KO cells and were selected for additional validation by immunocytochemical analysis. Primary dermal fibroblasts from a PD patient with homozygous *PINK1* Q456X loss-of-function mutation and matching WT cells were left untreated or were challenged with the mitochondrial depolarizer valinomycin and then stained with the top ten clonal supernatants (**Fig. S4A**). Most supernatants showed only minimal staining in untreated WT cells but detected a robust signal in stressed WT compared to *PINK1* mutant cells. Clones 11H2K3, 29H1K3, 29H2K2, and 30H3K1 had the most intense staining, while two clones showed no real signal increase above background levels (**Fig. S4B**).

Taken together and considering different biochemical and immunocytochemical validation criteria, cell supernatants from five recombinant p-S65-Ub clones (11H2K3, 19H1K1, 29H2K2, 30H3K1, and 41H1K3) were selected for subsequent affinity purification and further detailed characterization.

### Western blot assessment of the top affinity purified recombinant p-S65-Ub antibodies

Supernatant from the five top p-S65-Ub clones were next affinity-purified and first re-validated to confirm the previous outcome before expanding to additional analyses. Briefly, WT and *PINK1* KO HEK293E cells (**Fig. 2A**) as well as WT and *pink1* KO mouse embryonic fibroblasts (MEFs) (**Fig. 2B**) were treated with mitochondrial stressors and their lysates were analyzed by western blot. All recombinant affinity-purified p-S65-Ub antibodies were used in identical concentration, except the reference antibody, which was used five times diluted. All antibodies gave increased signal in WT but not the corresponding *PINK1*/*pink1* KO cells of human and mouse origin (**Fig. 2A, B**). In WT HEK293E cells, recombinant p-S65-Ub clones 29H2K2 and 30H3K1 were able to detect p-S65-Ub with highest intensity, closely followed by clone 11H2K3 and subsequently clones 19H1K1 and 41H1K3. Different from HEK293E cells, all clones except 41H1K3 performed comparably in treated WT MEFs. Overall, all five affinity-purified clones showed very little background signal in western blots of human and mouse *PINK1*/*pink1* KO cells treated with mitochondrial stressors, suggesting high specificity.

**Figure 2.**
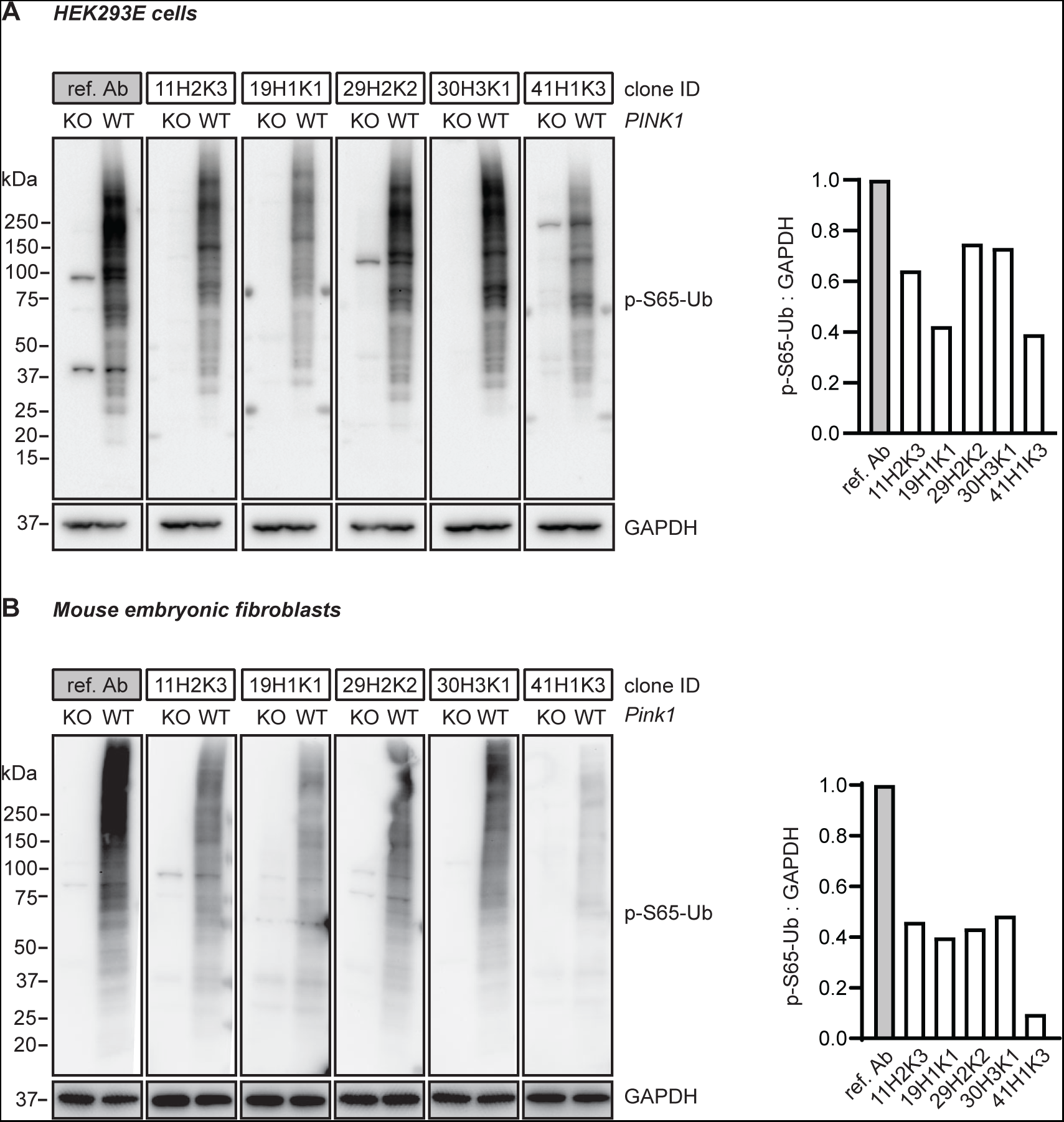
Western blot assessment of the top p-S65-Ub antibodies in cells. (**A**) HEK293E cells and (**B**) mouse embryonic fibroblasts with (WT) or without PINK1 expression (*PINK1* KO or *pink1* KO) were treated with mitochondrial depolarizer and analyzed by western blot. Representative western blot images are shown for the performance of the top five p-S65-Ub antibody clones in each cell type. Note that concentrations of the tested p-S65-Ub antibodies are five times higher compared to the reference antibody. GAPDH was used as loading control. A densitometric quantification of p-S65-Ub relative to GAPDH is shown alongside after normalization by the reference antibody set to 1. Ref. Ab - reference antibody, KO - knockout, WT - wild-type.

### Testing antibodies with recombinant p-S65-Ub chains using sandwich ELISAs

We next tested the purified recombinant p-S65-Ub antibodies in a sandwich ELISA format on a Meso Scale Discovery (MSD) platform. Rabbit p-S65-Ub antibodies were used as capturing agents and combined with a mouse total Ub antibody (P4D1) as detecting reagent as published previously [13]. We first tested recombinant p-S65-Ub tetramers with K48 or K63 chain linkage, the two most abundant Ub chain types in cells, and used their non-phosphorylated counterparts as negative controls and determined the Limit of Blank (LoB), the Limit of Detection (LoD), and the Limit of Quantification (LoQ) according to *Armbruster and Pry (2008)* [16] (**Table 1**; **Fig. 3A**). The LoB represents the relative noise or background, and was relatively high for clones 41H1K3, 11H2K3, and 29H2K2. The LoD was similar compared to the corresponding LoB and consequently also relatively high for the same clones mentioned (41H1K3, 11H2K3, and 29H2K2). Importantly, the LoQ for K48-linked p-S65-Ub tetramers was lowest for clone 19H1K1 at 100 fg/ml, which was even lower compared to the reference antibody.

**Figure 3.**
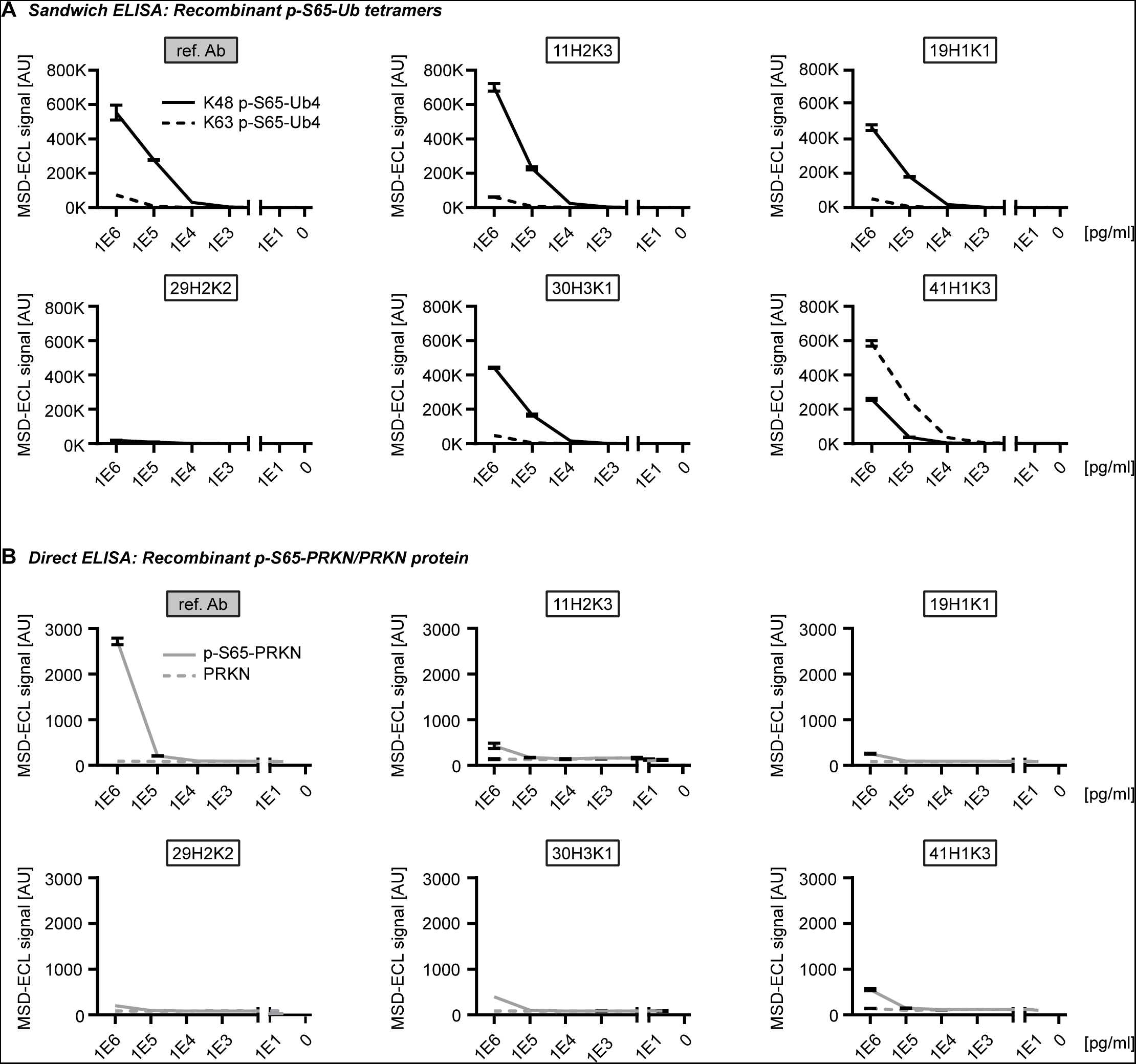
Assessing the top p-S65-Ub antibodies in ELISA using recombinant protein. (**A**) Sandwich ELISA results for the detection of recombinant K48 (solid line) and K63 (dash line) p- S65-Ub tetramers (Ub4) that were serially diluted to the range of 1-1,000,000 pg/ml. (**B**) Direct ELISA results for the detection of p-S65-PRKN and nonphosphorylated PRKN recombinant proteins that were serially diluted to the range of 1-1,000,000 pg/ml. N = 2. Data are shown as mean with SD.

**Table 1.**
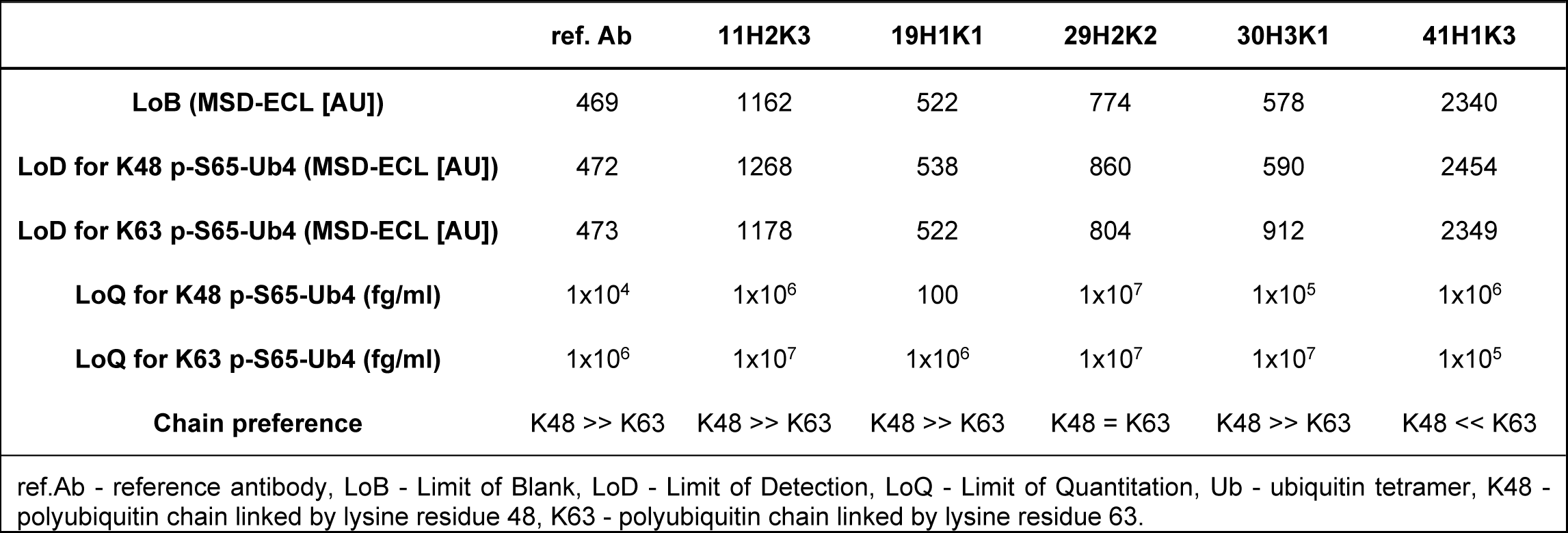
LoB, LoD, and LoQ of p-S65-Ub MSD ELISA with recombinant protein.

|  | ref. Ab | 11H2K3 | 19H1K1 | 29H2K2 | 30H3K1 | 41H1K3 |
| --- | --- | --- | --- | --- | --- | --- |
| **LoB (MSD-ECL [AU])** | 469 | 1162 | 522 | 774 | 578 | 2340 |
| **LoD for K48 p-S65-Ub4 (MSD-ECL [AU])** | 472 | 1268 | 538 | 860 | 590 | 2454 |
| **LoD for K63 p-S65-Ub4 (MSD-ECL [AU])** | 473 | 1178 | 522 | 804 | 912 | 2349 |
| **LoQ for K48 p-S65-Ub4 (fg/ml)** | 1x10<sup>4</sup> | 1x10<sup>6</sup> | 100 | 1x10<sup>7</sup> | 1x10<sup>5</sup> | 1x10<sup>6</sup> |
| **LoQ for K63 p-S65-Ub4 (fg/ml)** | 1x10<sup>6</sup> | 1x10<sup>7</sup> | 1x10<sup>6</sup> | 1x10<sup>7</sup> | 1x10<sup>7</sup> | 1x10<sup>5</sup> |
| **Chain preference** | K48 >> K63 | K48 >> K63 | K48 >> K63 | K48 = K63 | K48 >> K63 | K48 << K63 |

ref.Ab - reference antibody, LoB - Limit of Blank, LoD - Limit of Detection, LoQ - Limit of Quantitation, Ub - ubiquitin tetramer, K48 - polyubiquitin chain linked by lysine residue 48, K63 - polyubiquitin chain linked by lysine residue 63.

Of note, the LoQs for K63-linked p-S65-Ub tetramers were in most cases substantially higher compared to those for K48-linked p-S65-Ub tetramers, which is in line with previously published results [13]. Two exceptions were clones 41H1K3 and 29H2K2. Remarkably, clone 41H1K3 had a stronger preference towards Ub tetramers with K63-linkage relative to those with K48-linkage, which is reflected in a 10-fold lower LoQ for K63-linkage compared to Ub chains of the K48-linkage type. In contrast, the LoQ for both K48 and K63-linked p-S65-Ub were both similarly high (10 ng/ml each) for clone 29H2K2 suggesting that this antibody may not be the best option for ELISA-based detection (**Table 1**; **Fig. 3A**). The range of linearity in this assay differed between p-S65-Ub species used and was between 10 ng/ml and 1 μg/ml for K63 p-S65-Ub tetramers and between 1 ng/ml and 1 µg/ml for K48 p-S65-Ub tetramers. The coefficient of variation was excellent for all clones and matrices analyzed and was always well below 10% (data not shown).

Given the sequence similarity of phosphorylated epitopes around serine 65 in both Ub versus PRKN, we tested all purified recombinant antibodies towards phosphorylated and non-phosphorylated PRKN protein in a direct ELISA on an MSD platform (**Fig. 3B**). It is noteworthy that all antibodies including the reference antibody showed no cross-reactivity towards p-S65-PRKN or total PRKN except for the highest concentration tested (1 µg/ml). At this concentration, equimolar amounts of p-S65-Ub monomer resulted in almost 600-fold fold higher MSD-ECL signal for the reference antibody and between 2600 to 6600-fold higher MSD-ECL signal for all other recombinant p-S65-Ub clones. Lack of significant cross-reactivity with recombinant p-S65-PRKN protein was further confirmed by western blots (data not shown).

### ELISA-based p-S65-Ub detection in cells and in mouse and human brain lysates

We next tested the purified p-S65-Ub antibodies for ELISA using cell culture lysates (**Fig. 4A**). We used WT and *PINK1* KO HEK293E cells treated with CCCP for 24 h and compared to vehicle (DMSO)-treated WT cells. It is of note that all antibodies except clone 29H2K2 were able to show significant differences in p-S65-Ub levels at baseline condition between DMSO-treated WT and CCCP-treated *PINK1* KO cells. In line with previous results using recombinant p-S65-Ub chains, clone 19H1K1 and the reference p-S65-Ub antibody showed most sensitive p-S65-Ub detection and comparable results for both baseline (1.8- vs. 2.1-fold increases compared DMSO-treated WT to CCCP-treated *PINK1* KO cells) and upon stress (91- vs. 144-fold increases compared CCCP-treated WT over CCCP-treated *PINK1* KO cells) (**Fig. 4A**; **Table 2**). Interestingly, clone 41H1K3 that showed a higher sensitivity for recombinant K63- over K48-linked p-S65-Ub chains showed the lowest fold changes for p-S65-Ub detection in the cell samples, both at baseline and under mitochondrial stress (**Table 2**).

**Figure 4.**
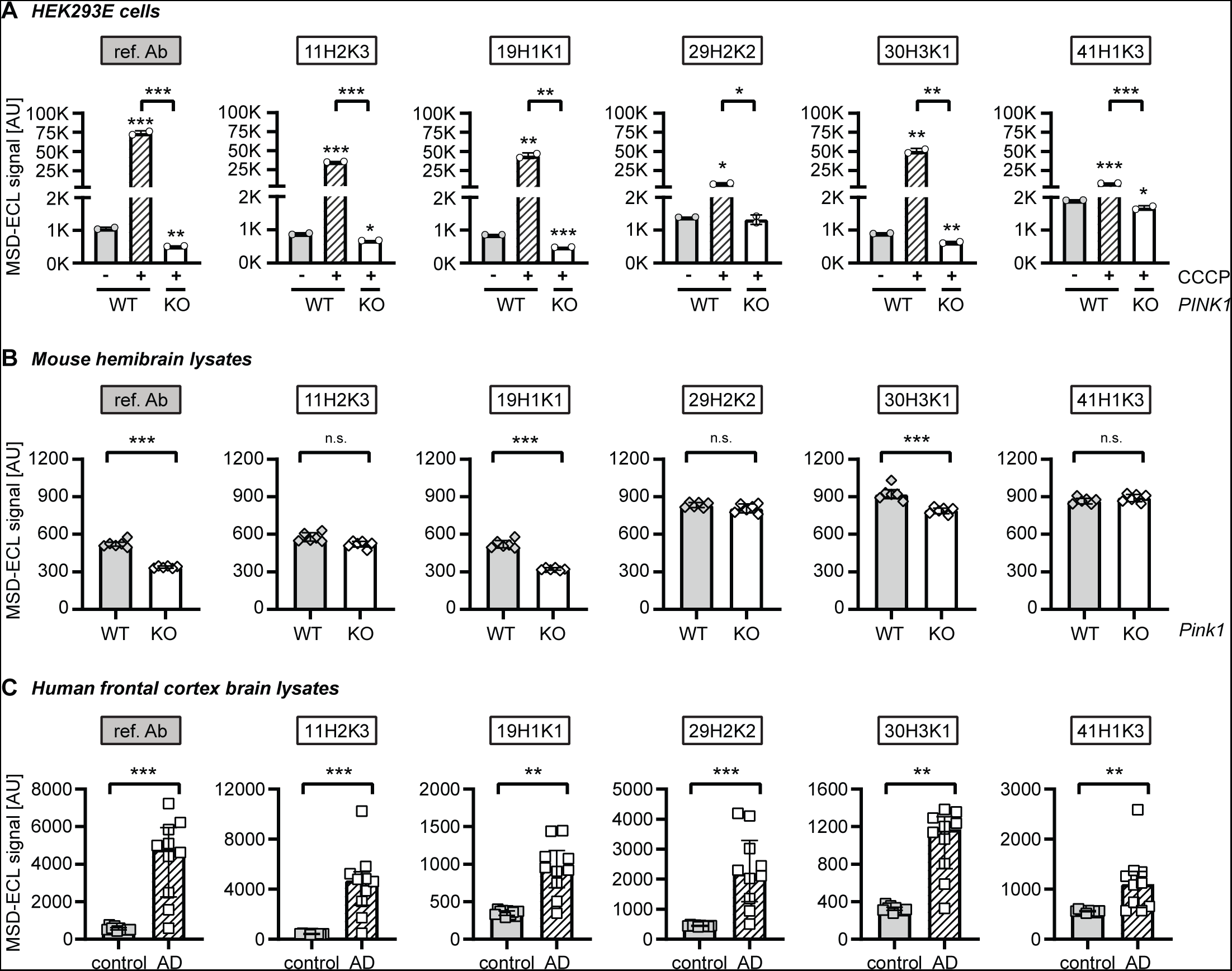
Assessing the top p-S65-Ub antibodies by sandwich ELISA using lysates from cells, mouse brain, and human brain. (**A**) ELISA detection of p-S65-Ub levels in cell lysates from CCCP or DMSO treated WT and *PINK1* KO HEK293E cells. One-way ANOVA. N = 2. Data are shown as mean with SEM. (**B**) ELISA detection of p-S65-Ub levels in hemibrain lysates from unstimulated WT and *pink1* KO mice. Unpaired t-test. N = 6 per group. Data are shown as mean with SEM. (**C**) ELISA detection of p-S65-Ub levels in human brain lysates from neurological normal controls and AD cases. Mann-Whitney U test. N = 10 per group. Data are shown as median with interquartile range. * p<0.05, ** p<0.005, *** p<0.0001. Asterisks (*) indicate the comparison to the untreated WT or the comparison between two groups under the brackets. Ref. Ab - reference antibody, WT - wild-type, KO - knockout, AD - Alzheimer disease, n.s. - non significant.

**Table 2.** Comparison of MSD ELISA detection of p-S65-Ub with different testing materials.

| Fold change in MSD ELISA detection | ref. Ab | 11H2K3 | 19H1K1 | 29H2K2 | 30H3K1 | 41H1K3 |
| --- | --- | --- | --- | --- | --- | --- |
| HEK293E cells baseline detection<br>(WT vs. *PINK1* KO) | 2.1 | 1.3 | 1.8 | 1.1 | 1.4 | 1.1 |
| HEK293E cells stressed condition<br>(treated WT vs. *PINK1* KO) | 143.5 | 52 | 91.2 | 5.6 | 80.2 | 4.8 |
| Mouse brain baseline detection<br>(WT vs. *pink1* KO) | 1.5 | 1.1 | 1.6 | 1 | 1.2 | 1 |
| Human brain disease condition<br>(control vs. AD) | 9.3 | 10.8 | 2.6 | 4.9 | 3.7 | 2 |

WT - wild-type, KO - knockout, AD - Alzheimer disease.

We then tested the performance of recombinant p-S65-Ub antibodies in mouse hemibrain lysates from unstimulated WT and *pink1* KO mice (**Fig. 4B**). As seen with recombinant Ub chains and in cells, clone 19H1K1 and the reference p-S65-Ub antibody again showed comparable results in mouse brains. Both antibodies were able to distinguish between the two genotypes with similar fold changes (1.6 vs. 1.5) (**Table 2**). A weaker detection of p-S65-Ub in unstimulated WT mice was found when using clone 30H3K1, while none of the remaining antibodies showed differences in p-S65-Ub levels between WT and *pink1^-/-^* mice (**Fig. 4B**; **Table 2**).

Finally, we tested the performance of purified p-S65-Ub clones in human autopsy brain lysates from the frontal cortex of Alzheimer disease (AD) cases compared to sex- and age-matched neurologically normal controls (**Fig. 4C**). Different from the results with mouse brain lysates, all p-S65-Ub clones detected significant increases of p-S65-Ub levels in AD cases compared to controls. The highest fold change between the two groups were obtained with clone 11H2K3 (10.8-fold) and the reference antibody (9.3-fold), followed by clones 29H2K2 and 30H3K1 and finally clones 19H1K1 and 41H1K3 (**Fig. 4C**; **Table 2**). It should be noted that the reference antibody, clones 19H1K1 and 30H3K1 showed the largest differences between control tissue and the buffer blank (data not shown), suggesting a greater assay window with potentially better detectability of low p-S65-Ub amounts.

### Characterization of p-S65-Ub clones via immunocytochemistry in PINK1 patient fibroblasts

To further confirm the specificity and sensitivity of the top five p-S65-Ub clones in other applications, we next validated them by immunocytochemistry in WT and homozygous *PINK1* loss-of-function mutation (*PINK1*^Q456X^) skin fibroblasts. For this, both WT and *PINK1* mutant fibroblasts were left untreated or were treated with the mitochondrial stressor valinomycin, stained with each of the antibodies using the same concentration and measured by high content imaging (**Fig. 5**). Both unstressed WT cells and stressed *PINK1* mutant cells were expected to show minimal p-S65-Ub signal and served as negative controls. Valinomycin-induced mitophagy led to a significant elevation of p-S65-Ub signal in WT fibroblasts as confirmed by the previously validated reference antibody [13] (**Fig. 5A**). All five clones also showed robust immunostaining of treated WT cells but only minimal immunopositive signals in untreated cells or treated *PINK1* mutant cells (**Fig. 5A**), indicating excellent specificity towards p-S65-Ub detection. In the treated WT cells, the highest signal was observed with antibody clones 30H3K1, 11H2K3, and 29H2K2. Clone 19H1K1 showed a moderate intensity, and 41H1K3 resulted in the lowest signal (**Fig. 5B**). The performance of the top three clones was comparable to the reference antibody. The specificity of the antibodies was further confirmed by the strong co-localization of p-S65-Ub positive signals with the mitochondrial marker HSP60 upon mitochondrial stress in WT cells (**Fig. 5C**).

**Figure 5.**
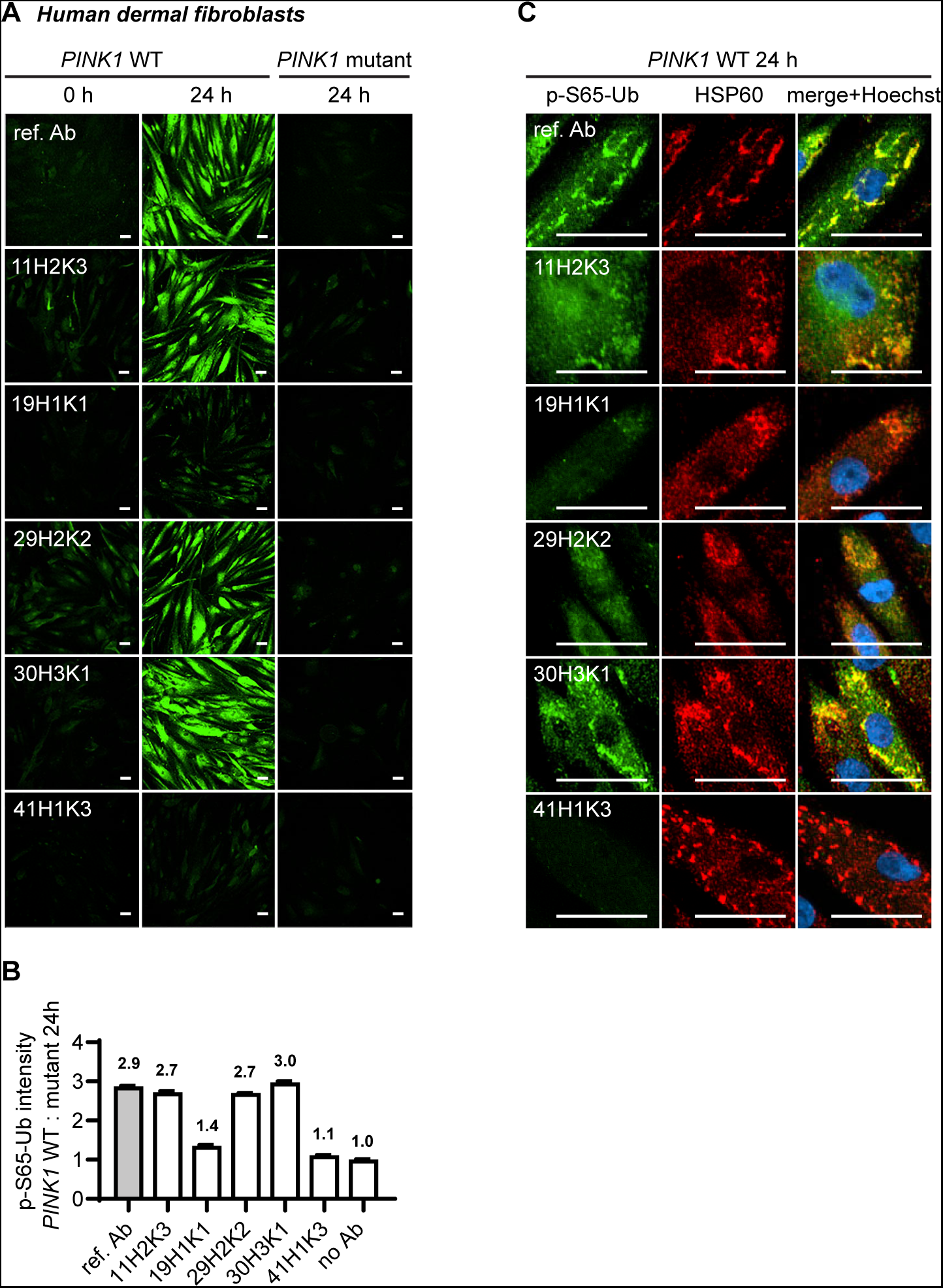
Comparison of the top p-S65-Ub antibodies using immunocytochemistry staining of human dermal fibroblasts. All five p-S65-Ub recombinant antibodies and the reference antibody were evaluated by immunocytochemistry in human primary dermal fibroblasts treated with 2 μM valinomycin for 0 or 24 hours. All antibodies were used at 1 μg/ml. (**A**) Representative images of p-S65-Ub immunoreactive signals (green) in fibroblasts carrying WT or homozygous *PINK1*^Q456X^ mutation. (**B**) Fluorescence intensities of each antibody was quantified by high content imaging and compared signals from treated WT to the signals from the corresponding *PINK1* mutant fibroblasts. Fold changes are labeled at top of each bar. Samples stained without primary antibody was used a negative control (no Ab). N = 2. Data are shown as mean with SEM. (**C**) Representative zoom-in images of p-S65-Ub and HSP60 immunoreactive signals in treated WT fibroblasts. 24 hours of valinomycin treatment induced mitochondrial aggregation (HSP60, red) and the accumulation of p-S65-Ub (green). Nuclei were stained with Hoechst (blue). Scale bar: 50 μm. Ref. Ab - reference antibody, WT - wild-type.

### Immunohistological assessment of p-S65-Ub clones in transgenic mouse and human AD brain

In addition to immunocytochemical staining in cultured fibroblasts, we next assessed p-S65-Ub labeling in the hippocampus of transgenic mice overexpressing human tau protein (rTg4510 mice). We have shown previously that p-S65-Ub is robustly induced in this tauopathy model and can reliably be detected by both immunohistochemical methods and MSD-ELISA [12,13,15]. Given that the reference antibody performs poorly in immunohistochemistry [13], the five p-S65-Ub recombinant antibodies were compared with a previously validated in-house p-S65-Ub antibody [11,14] (**Fig. 6**). Non-transgenic (nonTg) littermates were used as controls. Typical, punctate p-S65-Ub immunopositive structures (dark brown) were detected by all p-S65-Ub clones at identical concentrations with high specificity in sections from 9-10 months old transgenic but not nonTg mice. However, intensity levels were quite different for different recombinant clones and p-S65-Ub positive structures were most prominently detected with clone 41H1K3 followed by our in-house antibody [11,14,15], and minimal signal was found with the reference antibody (**Fig. 6**).

**Figure 6.**
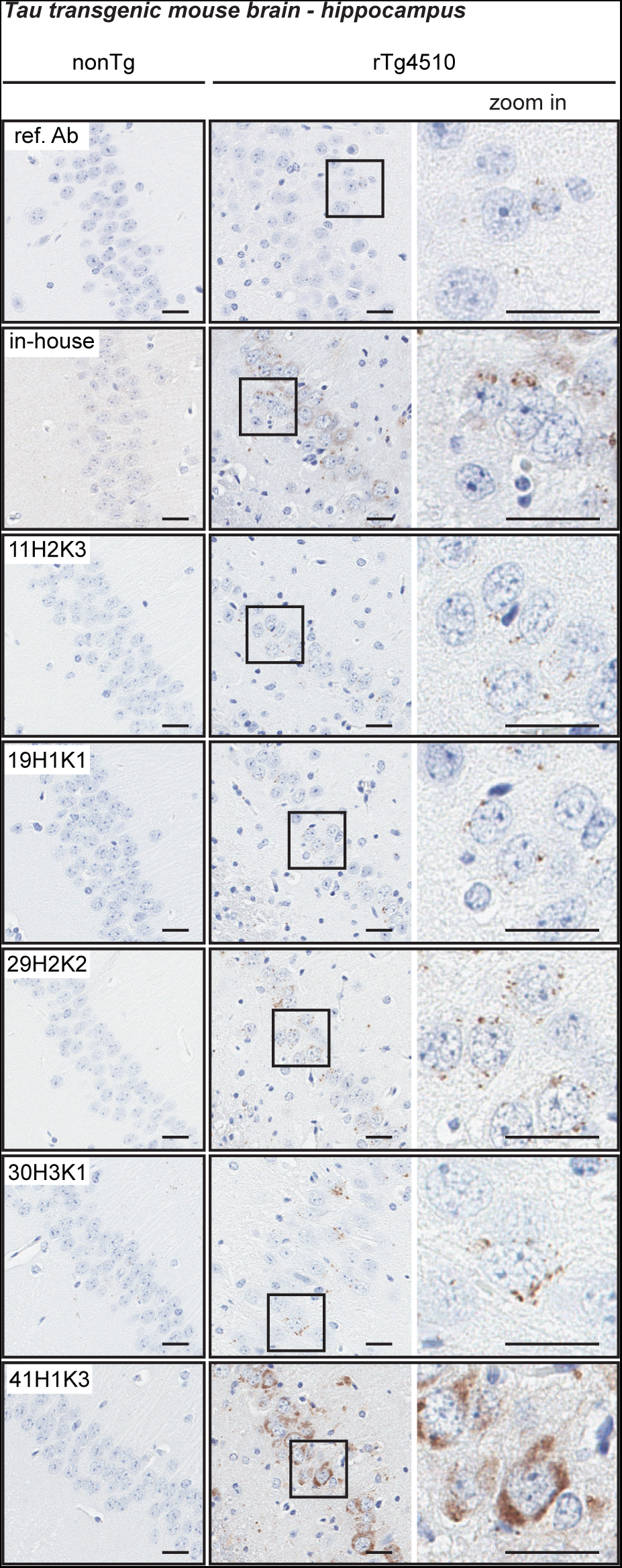
Characterization of the top p-S65-Ub antibodies by immunohistochemical staining of mouse brain. All five p-S65-Ub recombinant antibodies together with our in-house p-S65-Ub antibody and the reference antibody were evaluated by immunohistochemistry in brain sections from 9- to 10-month-old nonTg and rTg4510 mice. Same concentration (1.56 μg/ml) was used for all antibodies. Zoom-in images of the highlighted regions are shown to the right. Typical punctate p-S65-Ub immunopositive structures (dark brown) were detected by all clones but with different intensity. No signal was observed in nonTg mice. Scale bar: 50 μm. Ref. Ab - reference antibody, nonTg – non-transgenic.

Next, we tested the antibody clones on hippocampal regions of human AD brain (**Fig. 7**) [15]. Again, we used the same concentration (0.78 μg/ml) for all tested antibodies to directly compare the sensitivity. As previously reported, the in-house antibody detects small granular, cytoplasmic p-S65-Ub immunopositive structures in the AD hippocampus [15]. Similar p-S65-Ub structures were also observed when stained with clones 19H1K1, 29H2K2, and 41H1K3 with lower, similar, or higher signal intensity compared to the in-house antibody, respectively. Because of the sensitivity of 41H1K3 we diluted this antibody further and were still able to clearly label p- S65-Ub positive granules at a much lower concentration of 0.125 µg/ml (**Fig. 7**). In contrast to the abovementioned three clones, clone 11H2K3 showed undefined and rather diffuse cytoplasmic staining that has not been previously observed and only very minimal immunoreactive signal was found in clone 30H3K1-stained section.

**Figure 7.**
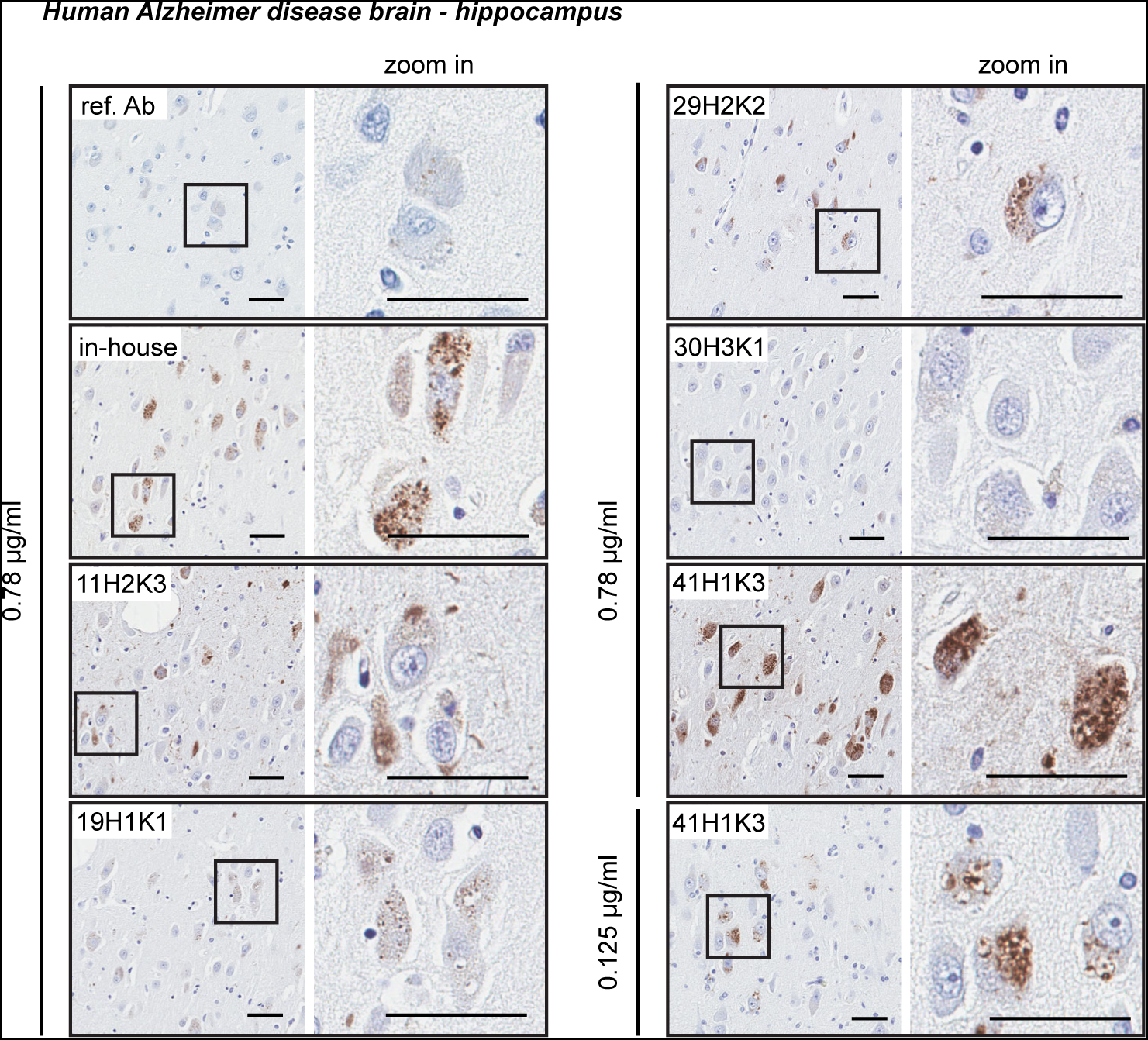
Characterization of the top p-S65-Ub antibodies by immunohistochemical staining of Alzheimer disease brain. All five p-S65-Ub recombinant antibodies together with our in-house p- S65-Ub antibody and the reference antibody were evaluated by immunohistochemistry of hippocampal sections from Alzheimer disease brain. Same concentration (0.78 μg/ml) was used for all antibodies. Zoom-in images of the highlighted regions are shown to the right. Typical punctate p-S65-Ub immunopositive structures (dark brown) were detected by most of the antibodies at different sensitivity. The superior sensitivity of 41H1K3 allowed detection of p-S65- Ub at a much lower concentration (0.125 μg/ml). Scale bar: 50 μm. Ref. Ab - reference antibody.

Taken together, selected clones showed great sensitivity for p-S65-Ub detection in immunostaining in fibroblasts (clones 11H2K3, 29H2K2, and 30H3K1) and brain tissue (clone 41H1K3). While similar characteristic morphologies were detected upon mitochondrial stress in cells, in tauopathy mouse brain, and in human disease autopsy brain, the clones that worked the best for immunostaining in cells and tissue did not appear to overlap.

## DISCUSSION

We here developed and thoroughly characterized a new series of recombinant, rabbit monoclonal antibodies targeting the mitophagy tag p-S65-Ub. Starting from an immunization of four animals, we screened over 200 B cell lines by direct ELISA and cloned and sequenced 47 recombinant antibodies that were further evaluated. The top five clones were affinity purified and their sensitivity and specificity were validated using a range of biochemical and imaging techniques with recombinant Ub chains, gene-edited cells, and patient fibroblasts as well as mouse and human brain tissue. We compared their respective signal-to-noise ratios in samples with diverse complexity and levels of the mitophagy tag at baseline and upon acute, mitochondrial stress or in chronic, diseased conditions, and across applications. The top performing antibodies were selected for each category and are now available to the community, expanding the toolkit to monitor PINK1-PRKN signaling in different sample types and with different methods (see Table 3 for a summary).

**Table 3.** Top recombinant p-S65-Ub antibodies for different applications.

| Matrix | Cells |  |  | Brain tissue |  |
| --- | --- | --- | --- | --- | --- |
|  | WB | ELISA | ICC | ELISA | IHC |
| human | 30H3K1* | 30H3K1* | 30H3K1* | 30H3K1* | 41H1K3* |
|  | 19H1K1 | 19H1K1 |  | 19H1K1 |  |
|  | 41H1K3 |  |  |  |  |
| mouse | 30H3K1* | ND | ND | 19H1K1* | 41H1K3* |
|  | 19H1K1 |  |  | 30H3K1 |  |

\* = denotes best clone for the application in that sample type. WB - western blot, ICC - immunocytochemistry, IHC - immunohistochemistry, ND - not defined.

Since its discovery [17–19], p-S65-Ub has been recognized as a highly specific and quantitative mitochondrial damage marker of disease relevance [14,20,21]. By now p-S65-Ub has been employed for larger-scale, functional and genetic screening of mitophagy modifiers in cell culture [22,23] and in human autopsy brain [24]. p-S65-Ub can also be detected in clinical samples and may be useful as a biomarker of mitochondrial health and function [13,25]. Yet, physiological levels of p-S65-Ub can vary between different cells and tissues and the normal range of the p- S65-Ub signal needs to be established for each case and sample type. While p-S65-Ub is strongly elevated with stress, age, and various disease conditions in cell and animal models and in humans [11–13,15,26], the underlying causes may differ. It is of note that while p-S65-Ub is a direct result of PINK1 and PRKN enzymatic activities, this mitophagy label is only transitory and highly dynamic in nature as damaged labeled mitochondria are rapidly degraded under healthy, physiological conditions. Greater levels of p-S65-Ub can result from primarily increased mitochondrial stress and damage or from an impaired turnover and reduced flux through the autophagy-lysosome system. Monitoring p-S65-Ub levels over time may allow to better determine mitophagic flux and could help discriminate between both scenarios. Therefore, it may be useful not only as a diagnostic and prognostic tool, but also as pharmacodynamic and therapeutic biomarker.

Although mitochondrial proteins are the primary targets of p-S65-Ub, the exact nature and composition of the phosphorylated Ub chains is not well understood but could have important implications for outcomes and detection of the signal. A preference for phosphorylation of multi-mono-Ub moieties or short poly-Ub chains has been suggested in cell lines [27], but an increase particularly in K63-linked poly-Ub chains has been noted on mitochondria and appears to be the preferred mitophagy signal in neurons [28,29]. The temporal order of events as well as a certain stoichiometry of the p-S65-Ub signal could also be important for proper induction and flux of mitophagy. As such p-S65-Ub specific antibodies that are also able to better discriminate between different Ub linkages might be particularly useful to further resolve the mitophagy signal *in vivo*. In that regard it is worth noting that some of the clones developed herein seemed to have somewhat distinct chain linkage preferences. While most antibodies to date including clones 11H2K3, 19H1K1, and 30H3K1, seem to have a higher affinity towards recombinant K48-linked p-S65-Ub chains, clone 41H1K3 preferred phosphorylated K63-linked chains. Clone 29H2K2 seemed to recognize both types equally well. Certainly, much more work is needed to fully characterize the binding profiles and determine selectivity and affinity of the antibody clones, but a toolkit consisting of clones with differential binding abilities combined with sophisticated mass spectrometry approaches [30] may provide a unique opportunity to decode the complexity of the p-S65-Ub signal in health and disease.

We interrogated the top five antibodies across a range of methods using cell and brain tissue samples of mouse and human origin alongside with appropriate controls for each setting. It is noteworthy that none of the top five clones performed best in all tested applications or with all sample types. Instead, 30H3K1, 19H1K1, and 41H1K3 were selected as the best clones for different applications and are currently in production for public access (see details in Materials and Methods). Clone 30H3K1 convinced particularly through its performance in western blots and immunocytochemistry, clone 41H1K3 showed the highest sensitivity for immunohistochemistry in mouse and human brain tissue, and clone 19H1K1 worked best in sandwich ELISAs using cell culture and (mouse) brain lysates. Although other clones such as 11H2K3 also performed quite well in MSD-ELISA and perhaps captured even better the peak intensities under more severe conditions (e.g., human AD brain tissue), clone 19H1K1 had a bigger assay window towards the lower end of the signal. It appeared most useful to also detect the basal, physiological p-S65-Ub levels and subtle changes within cells, normal mouse brain, and human control brain. Whether differences in the sample matrix, changes in the Ub chain composition, or linkage preferences influenced the clones’ overall performance in different applications or sample types remains to be determined. However, testing of the clones should be expanded to clinical biofluids in order to determine their full potential for also ultrasensitive detection of p-S65-Ub.

Here we capitalized on recent technological developments in the field and generated first recombinant, rabbit monoclonal antibodies against the mitophagy tag p-S65-Ub. Compared to conventional monoclonal antibodies derived from hybridoma cells lines, recombinant antibodies are not susceptible to genetic drift, gene loss, or mutations. As such recombinant antibodies are more stable and better scalable with minimal variability in specificity and excellent batch-to-batch reproducibility. Antibodies may also be recloned into backbones from a different species or conjugated with other epitope tags or direct site-specific labeling. Overall, this could generate more flexibility and enhance their versatility across applications where the host species may be limiting their utility [31]. Given the known sequences of the heavy and light chain variable regions, recombinant antibodies can be further engineered. For instance, the variable regions could be modified to further modulate the affinity to different Ub linkages, making them more selective for a certain Ub chain type. It was of interest that clones 11H2K3 and 30H3K1 were located on one end, while clone 41H1K3 was located on the opposite end on the phylogenetic tree. It is however not clear whether there is a correlation between the distance and their binding preferences with regards to Ub chain linkages.

In summary, we here introduce and characterize novel recombinant p-S65-Ub antibodies that can now be further exploited by the field to better understand mechanisms and nature of the PINK1-PRKN mediated p-S65-Ub signaling in a range of applications as well as its underlying implication in various disease-associated conditions. Our study also highlights the biomarker potential of the mitophagy tag in various clinical and pathological specimens as a diagnostic and prognostic tool or for therapeutic assessment.

## MATERIAL AND METHODS

### Peptide synthesis, rabbit immunization, and polyclonal antibody production

Synthetic peptides and the polyclonal antibodies were manufactured by 21^st^ Century Biochemicals, Marlborough, MA. Briefly, peptides were manufactured by Fmoc-based solid phase peptide synthesis, HPLC purified to >90%, and conjugated via the cysteine thiol to a mixture of carrier proteins. Each peptide was covalently attached to Optimized Affinity Resin (21^st^ Century Biochemicals) containing an extended spacer arm and a thiol-reactive moiety. All rabbit animal work was conducted at Pacific Immunology (San Diego, CA; NIH and USDA compliant). After an initial rabbit immunization and multiple boosts, the serum was collected at specific intervals and analyzed by dot blots at 21^st^ Century Biochemicals after which appropriate production bleeds were pooled. The serum then underwent two purification processes. Antibodies were first recovered by affinity purification using p-S65-Ub peptides covalently attached to Optimized Affinity Resin through a cysteine residue added to either the N- or C-terminus. A portion of the resulting antibodies were further subjected to an additional immunodepletion step with both the nonphosphorylated Ub peptides (Ub Ser-OH N-/C-terminus) and PRKN peptides (p-S65-PRKN control and Ser-OH65-PRKN control). Peptide affinity columns were incubated with serum, washed multiple times including a 0.5 M NaCl wash in phosphate buffer, and the antibodies eluted under acidic conditions. After dialysis against PBS, the antibodies were stored in PBS with 50% glycerol.

### PBMC isolation, B cell selection, screening, and recombinant antibody generation

Following the final boost, 30 ml of rabbit peripheral blood was collected for PBMC isolation and sent to ImmunoPrecise Antibodies Ltd. In short, the whole blood from each animal was diluted one to one in serum free medium and subjected to a density separation medium, Lympholyte-Mammal (Cedarlane Laboratories, L5110). Following the manufacturer’s protocol, the enriched PBMCs at the middle layer were carefully removed and washed twice in 1% BSA (VWR, 9048- 45-8) in DMEM (Corning, 15-018-CV). The isolated PBMCs were then subjected to an antigen specific enrichment process as part of the ImmunoPrecise Antibodies Ltd. Proprietary B Cell Select platform and were diluted in growth medium for single B cell cultivation. On day 7, the supernatant of each single B cell well was screened for antigen specificity through direct ELISA for targeting full-length monomeric p-S65-Ub protein (Boston Biochem, U-102), free p-S65-Ub peptides 1 and 2 and BSA-conjugated p-S65-Ub peptides 1 and 2 (provided by 21^st^ Century Biochemicals). Non-phosphorylated Ub protein (Boston Biochem, U-100H), free non-phosphorylated, hydroxylated Ub peptides 1 and 2, free p-S65-PRKN peptide, and free hydroxylated p-S65-PRKN peptide (provided by 21^st^ Century Biochemicals) were used as negative controls.

The top 48 performing B cells from the screening were assessed and selected for recombinant expression. Briefly, mRNA was isolated (Dynabeads, Invitrogen, 61006) from lysed B cell cultures and RT-PCR was performed for the conversion to cDNA, followed by amplification of targeted heavy and light chains using specific primer sets. Variable regions of G1 and kappa chains were separately ligated into expression vectors containing respective constant regions. Plasmids encoding both the antibody heavy chain and light chain, housed within an expression vector driven by the CMV promoter were co-transfected into HEK293 cells, which had been adapted for suspension culture in a serum-free medium. Following transfection using the polyethylenimine-mediated gene delivery method, the cells were allowed to incubate for a further six days, resulting in 47 single clones that were producing recombinant antibodies. Variable region sequences of antigen specific antibodies were determined by analysis of samples submitted to Psomagen, MD, USA (**Table S1; Fig. S2**).

Sequences for the top five recombinant antibodies were sent to Abcam for production of purified recombinant antibodies and commercial distribution. The five top performing antibodies produced in the supernatant were captured by Protein A ligands, specifically binding to the constant region (Fc) of the IgG antibodies (CH2-CH3 region) as the final purification step. The three best clones 30H3K1 (Abcam, ab309155, 30H3/30K1), 19H1K1 (Abcam, ab314901, 19H1/19K1), and 41H1K3 (Abcam, ab314903, 41H1/41K3) for different applications as recommended in **Table 3** are publicly available (https://www.michaeljfox.org/research-tools-catalog).

### Dot blots using peptides

In peptide dot blots, phosphorylated and non-phosphorylated Ub 13-mers with central Ser65 respective PRKN 12mers (Ser65 in 6^th^ position) were used in decrements of 5 (left to right) starting with 25 µM, 5 µM, 1 µM and 0.2 µM concentrations on 0.2 µm Nitrocellulose membrane (BioRad, 10484059). Ub peptides 1 and 2 had identical peptide sequences but different N- and C-terminal linker residues e.g., 6-Aminohexanoic Acid. The membrane was placed between 2 sheets of filter paper (Whatman, 1001917) and dried overnight. The next morning the membrane was rinsed in Tris-buffered saline (TBS; 50 mM Tris [Millipore, G48311], 150 mM NaCl [FisherScientific, BP358], pH 7.4) for ∼10 seconds, blocked in 5% skim milk (Sysco, 5398953) in TBS with 0.1% Tween (TBS-T) for 30 minutes at room temperature. Primary antibodies were then added in a concentration of 1 μg/ml in 5% BSA (Boston BioProducts, P-753) in TBS-T and incubated for 2 hours at room temperature. Membranes were subsequently washed four times for 5 minutes in TBS-T before secondary antibody (Jackson ImmunoResearch, 711-035-152, 1:15,000) was added for 1 hour at room temperature in 5% skim milk. Membranes were washed five times for 10 minutes each in TBS-T and detected with Immobilon Western Chemiluminescent HRP Substrate (Millipore Sigma, WBKLS0500) on Pro Signal Blotting film (Genesee Scientific, 30-810L).

### Generation of MEFs

The *pink1* KO mice was obtained from Jackson Laboratories (RRID:IMSR_JAX:017946). Experiments were conducted in accordance with the Animal Scientific Procedures Act (1986) and with the Directive 2010/63/EU of the European Parliament and of the Council on the protection of animals used for scientific purposes (2010, no. 63). MEFs were generated using E13.5 embryos. Embryos were separated from placenta and individually placed into a single dish with cold DPBS, no calcium, no magnesium (Gibco, 14190094). After dissection of the head and all internal organs, embryo bodies were chopped with a spring scissor to mince the tissue of 1-2 mm in Petri dish. Small piece of tail was collected for genotyping. Tissues were digested for 15 min at 37 °C with digestion medium [0.025% Trypsin-EDTA (Gibco, 25300054); 0.125mg/mL DNase I (Merck, 11284932001) in HBSS (Gibco, 14025050)]. After enzymatic digestion, tissue dissociation was completed by pipette trituration and trypsin neutralised with 5 ml of culturing media [DMEM (Gibco, 11960-085); 20% heat-inactivated Fetal Bovine Serum (Gibco, 10500064); 1% Penicillin-Streptomycin (Gibco, 15140122), 1% L-Glutamine (Gibco, 25030024); 1X Non-essential Amino acid (Gibco, 11140-035), 1X Sodium pyruvate (Gibco, 11360-039)]. Then, dissociated tissues were centrifuged at 1200 rpm for 5 min and resuspend in 5 ml culturing media. Cell suspension was filtered using a 70 μm cell strainer in order to remove any undigested tissue pieces or aggregates from the cells. Cells were seeded in 10 cm dishes, containing 10 ml of pre-warmed culturing media, and cultured at 37°C in a humidified incubator with 5% CO_2_. Media was changed every 5 days and cells split at 90% confluence. A detailed protocol describing the preparation of mouse embryonic fibroblasts has been reported [32].

### Cell Culture and treatment

WT and *PINK1* KO HEK293E cells were previously described [13] and were cultured in Dulbecco’s modified Eagle medium (Thermo Fisher, 11965118) supplemented with 10% fetal bovine serum (Neuromics, FBS001800112). Primary human dermal fibroblasts were collected from a homozygous *PINK1*^Q456X^ carrier and a healthy relative to serve as negative control [33]. All fibroblasts were cultured in Dulbecco’s modified Eagle medium supplemented with 10% fetal bovine serum, 1% PenStrep (Thermo Fisher, 15140122), and 1% non-essential amino acids (Thermo Fisher, 11140050). WT and *pink1* KO MEF cells have been described previously [28]. MEFs were grown in Dulbecco’s modified Eagle medium supplemented with 20% fetal bovine serum, sodium pyruvate (Fisher Scientific, 11360-070), non-essential amino acids, and PenStrep. All cells were grown at 37°C, 5% CO_2_:air in humidified atmosphere. To induce mitophagy, HEK293E cells, MEFs, and human dermal fibroblasts were treated with 20 μM CCCP (Sigma-Aldrich, C2759), 5 μM valinomycin (Cayman Chemicals, 10009152), or 2 μM valinomycin, respectively, for 24 hours. DMSO (Sigma-Aldrich, D4540) was used as vehicle treatment.

### Western blotting

HEK293E cells and MEFs were lysed in RIPA buffer (50 mM Tris, pH 8.0, 150 mM NaCl, 1% NP-40 [United States Biological, 19628], 0.5% deoxycholate [Sigma-Aldrich, D6750], 0.1% SDS [Bio-Rad, 1610302]) containing protease inhibitor cocktail and phosphatase inhibitors (Sigma-Aldrich, 11697498001 and 04906837001). Cell lysates were then cleared for 15 minutes, 4°C at 20817 x g and protein concentrations were determined by BCA assay (Thermo Fisher, 23225). Cell lysates (10 μg for HEK293E cells or 15 μg for MEFs) were diluted in Laemmli buffer (62.5 mM Tris, pH 6.8, 1.5% SDS, 8.33% glycerol [Fisher Scientific, BP2291], 1.5% β-mercaptoethanol [Sigma-Aldrich, M3148], 0.005% bromophenol blue [Sigma-Aldrich, B5525]) and subjected to SDS-PAGE using 8-16% Tris-glycine gels (Invitrogen, EC60485BOX). Proteins were transferred onto polyvinylidene fluoride membranes (Millipore, Immobilon PSQ, ISEQ00010 or PVH00010) and the membranes were then blocked for 1 hour in 5% non-fat milk in TBS-T and incubated in primary antibodies overnight in 5% BSA in TBS-T at 4°C. 1 μg/ml of recombinant antibodies and 0.2 μg/ml of the reference antibody (Cell Signaling Technology, 62802) were used. GAPDH (Meridian Life Science, H86504M, 0.025 µg/ml) and vinculin/VCL (Sigma, V9131, 0.016 µg/ml) were used as loading controls. The following day, membranes were subjected to 1 hour incubation with secondary antibodies (Jackson ImmunoResearch, 711-035-152 and 715-035-150) at room temperature. Proteins were visualized using Immobilon Western Chemiluminescent HRP Substrate (Millipore Sigma, WBKLS0500) on Pro Signal Blotting film (Genesee Scientific, 30-810L) or on a Chemidoc MP imaging system (Bio-Rad, Hercules, CA, USA). Quantification of western blots was performed using Image Studio Lite software (Li-COR Biosciences, Version 5.2). Intensity levels per area were background subtracted and normalized to the loading controls.

### Tissue collection and preparation of mouse and human brain lysates

A total of 12 WT and *pink1* KO mice (N = 3 per sex/genotype) at age of three months were anesthetized and intracardially perfused with cold PBS (Invitrogen, 14190-235). Brains were removed and immediately flash-frozen in liquid nitrogen. All animal procedures were in accordance with the ethical standards established by Mayo Clinic and approved by the Institutional Animal Care and Use Committee at Mayo Clinic. Additionally, frozen human frontal cortex from ten AD and ten age- and sex-matched neurologically normal control cases were obtained from the Mayo Clinic Florida Brain Bank. All samples are from autopsies performed after approval by the legal next-of-kin. Research on de-identified postmortem brain tissue is considered exempt from human subjects’ regulations by the Mayo Clinic Institutional Review Board.

For protein extraction, frozen brain tissues were homogenized in 5x volumes of ice-cold RIPA buffer supplemented with protease and phosphatase inhibitors using glass Dounce tissue grinders (Fisher Scientific, K885300-0002). After 30 minutes incubation on ice, brain lysates were cleared by two times centrifugation at 20,000 x g, 4°C for 15 minutes. Resulting supernatant was collected as final lysates and protein concentrations were determined by BCA assay.

### Sandwich MSD ELISA using recombinant protein, cell lysates, and tissue lysates

Similar to the p-S65-Ub sandwich MSD ELISA that has been described previously [13], the top five recombinant p-S65-Ub antibodies or the reference antibody (Cell Signaling Technology, 62802, E2J6T) were used as capturing agents. Detection was performed with a mouse Ub antibody (ThermoFisher, 14-6078-37, P4D1) followed by a Sulfo-tag labeled anti-mouse antibody (MSD, R32AC-1). All capturing antibodies were used at a concentration of 1 µg/ml in 200 mM sodium carbonate buffer pH 9.7 and coated overnight at 4°C with 30 µl per well in 96-well MSD plate (MSD, L15XA-3). The next morning MSD plates were washed two times with TBS-T and incubated with 300 µl filtered blocking buffer (150 mM Tris, pH 7.4, 150 mM NaCl, 0.1% Tween-20, 1% BSA) for 1 hour at 22°C. Antigen incubation was performed for 2 hours at room temperature at 600 rpm on a microplate mixer (USA Scientific, 8182-2019). After washing, 30 µl of detecting mouse Ub antibody (ThermoFisher, 14-6078-37, P4D1, 5 µg/ml) were added to each well and incubated again for 2 hours. 50 µl Sulfo-tag labeled goat anti-mouse antibody (MSD, R32AC-1, 1:500) was next added and incubated for 1 hour. The plates were then read on a MESO QuickPlex SQ 120 reader (Meso Scale Diagnostics, Rockville, MD, USA) after addition of 150 µl MSD GOLD Read Buffer (MSD, R92TG-2).

For detection of recombinant proteins, K48 (Boston Biochem, UC-250) and K63 (Boston Biochem, UC-350) p-S65-Ub tetramers were serially diluted to a final concentration of 10 fg/ml to 1 µg/ml. Their nonphosphorylated counterparts (Boston Biochem, UC-210b and UC-310b) were used as negative controls. For detection in HEK293E cell lysates, 10 µg of protein was used with CCCP-treated *PINK1* KO cells serving as negative control and CCCP-treated WT cells serving as positive control. For detection in mouse and human brain lysates, 30 µg of protein was used. All samples were run in duplicates and diluted in blocking buffer.

The LoB, LoD, and LoQ were calculated using MSD-ECL values from recombinant p-S65-Ub chains and according to *Armbruster and Pry* [16]. In brief, the LoBs were determined using the formula LoB = mean _blank_ + 1.645(SD _blank_) where blocking buffer served as blank. The LoD was determined using serial dilutions of p-S65-Ub analytes and calculated with the following formula: LoD = LoB + 1.645(SD _low concentration sample_). The LoQ was determined as the lowest concentration of analyte that could be measured with a predefined goal for imprecision (here under 10%) or is clearly distinguishable from the LoB with more than 3 times the standard deviation.

### Direct MSD ELISA using recombinant protein

Recombinant p-S65-PRKN (Boston Biochem, E3-166), non-phosphorylated PRKN (Boston Biochem, E3-160) and p-S65-Ub monomer (Boston Biochem, U102) were initially dissolved in RIPA buffer (stock concentration of 1 µg/µl), aliquoted in 1 µl aliquots, flash-frozen in liquid nitrogen and stored at −80⁰C. Recombinant PRKN and p-S65-PRKN aliquots were serially (10-fold) diluted in 200 mM sodium carbonate buffer pH 9.7 with a starting concentration of 1 µg/ml towards 10 fg/ml and used to coat the MSD plates overnight at 4⁰C without shaking with 50 µl/well. Recombinant p-S65-Ub monomer (positive control) was used in equimolar concentration relative to the highest p-S65-PRKN concentration of 1 µg/ml. On the next morning, plates were washed twice with TBS-T and incubated for 1 hour with blocking buffer without shaking at 22°C. Recombinant p-S65-Ub antibodies and the reference antibody in blocking buffer were subsequently incubated at a concentration of 1 µg/ml with 30 µl/well for 2 hours at room temperature at 600 rpm on a microplate mixer. 50 µl Sulfo-tag labeled goat anti-rabbit Ab (MSD, R32AB-1, 1:500) was next added and incubated for 1 hour. The plates were then read on a MESO QuickPlex SQ 120 reader after addition of 150 µl MSD GOLD Read Buffer.

### Immunofluorescence staining of fibroblasts followed by high content imaging

To evaluate top clones for immunocytochemistry and quantify p-S65-Ub levels using automated high-content imaging, WT and homozygous *PINK1*^Q456X^ mutant fibroblasts were seeded in 96-well imaging plates (Fisher Scientific, 08772225) and allowed to attach overnight. Cells were treated for 2 µM valinomycin (Enzo Life Science, BML-KC140-0025) or vehicle DMSO (Sigma-Aldrich, D4540) for 24 hours and then fixed in 4% paraformaldehyde (Sigma-Aldrich, 441244) after one wash with PBS. Fibroblasts were immunostained with all clones or reference antibody against p-S65-Ub (Cell Signaling Technology, 62802, E2J6T) at 1 µg/ml and with HSP60 (arigo Biolaboratories, ARG10757, 1:2,000) followed by incubation with secondary antibody (Invitrogen, A-11034 and A-11041; 1:1,000) and Hoechst 33342 (Invitrogen, H21492; 1:5,000). Plates were imaged using an Operetta CLS High Content imaging system (Perkin Elmer, Waltham, MA, USA) with a 20x water immersion objective using a 2x2 montage (no gaps) per well. Raw images were processed using the built-in Harmony analysis software to calculate the average cytoplasmic p-S65-Ub intensity per cell in each well.

### Processing of fixed mouse and human tissue and immunohistochemistry

Without doxycycline administration, rTg4510 mice used here constitutively express high levels of human P301L tau in the hippocampus. Brain tissue from one nonTg and one rTg4510 transgenic mice at 9 months of age were used for immunohistochemistry. All procedures were in accordance with the ethical standards established by Mayo Clinic and approved by the Institutional Animal Care and Use Committee at Mayo Clinic. Additionally, autopsy brain from a single AD case was retrieved for immunohistochemistry from the Mayo Clinic Florida Brain Bank.

Paraffin embedded hippocampal tissue was cut into 5 µm consecutive sections and allowed to dry overnight at 60°C. Slides were deparaffinized and rehydrated, followed by antigen retrieval in steaming deionized water for 30 minutes. After blocking with 0.03% hydrogen peroxide (Sigma, 216763) and 5% normal goat serum (Invitrogen, 16210072), sections were incubated with all five clones, reference p-S65-Ub antibody (Cell Signaling Technology, 62802, E2J6T), or in-house p-S65-Ub antibody at 1.56 µg/ml (in mouse brain), 0.78 µg/ml (in human brain), or 0.125 µg/ml (only for clone 41H1K3 in human brain), followed by rabbit-labeled polymer HRP (Agilent, K4011) at room temperature. Peroxidase labeling was visualized with the chromogen solution 3,3’-diaminobenzidine (Agilent, K4011). The sections were then counterstained with Lerner 1 hematoxylin (Fisher Scientific, CS400-1D) and coverslipped with Cytoseal mounting medium (Thermo Scientific, 8310). After immunohistochemical staining, all sections were imaged with an Aperio AT2 digital pathology scanner at 40x magnification (Leica Biosystems, Wetzlar, Germany).

### Statistical analysis

Data analysis was performed in GraphPad Prism version 9 software. One-way ANOVA followed by Tukey’s multiple comparison test was used for the HEK293E cell ELISA analysis. Unpaired t-test and Mann-Whitney U test were used for mouse and human brain ELISA analysis, respectively.

## ACKNOWLEDGEMENTS

We thank The Michael J. Fox Foundation for Parkinson’s Research for their guidance and leadership in this collaborative effort. We are also grateful to the patients and their families who made this study possible. We thank the Center for Regenerative Medicine and the Neuroregeneration Laboratory for biobanking of patient cells at Mayo Clinic Florida.

## ABBREVIATIONS

AD: Alzheimer disease
CCCP: carbonyl cyanide 3-chlorophenylhydrazone
ELISA: enzyme-linked immunosorbent assay
HEK293E cell: human embryonic kidney E cell
KO: knockout
LoB: Limit of Blank
LoD: Limit of Detection
LoQ: Limit of Quantification
MEF: mouse embryonic fibroblast
nonTg: non-transgenic
PBMC: peripheral blood mononuclear cell
PD: Parkinson disease
p-S65-PRKN: phosphorylated PRKN at serine 65
p-S65-Ub: phosphorylated Ub at serine 65
Ub: Ubiquitin
VCL: vinculin
WT: wild-type

