## Supplemental Materials for "Development and characterization of phospho-ubiquitin antibodies to monitor PINK1-PRKN signaling in cells and tissue"

**Running title:** Development of recombinant phospho-ubiquitin antibodies

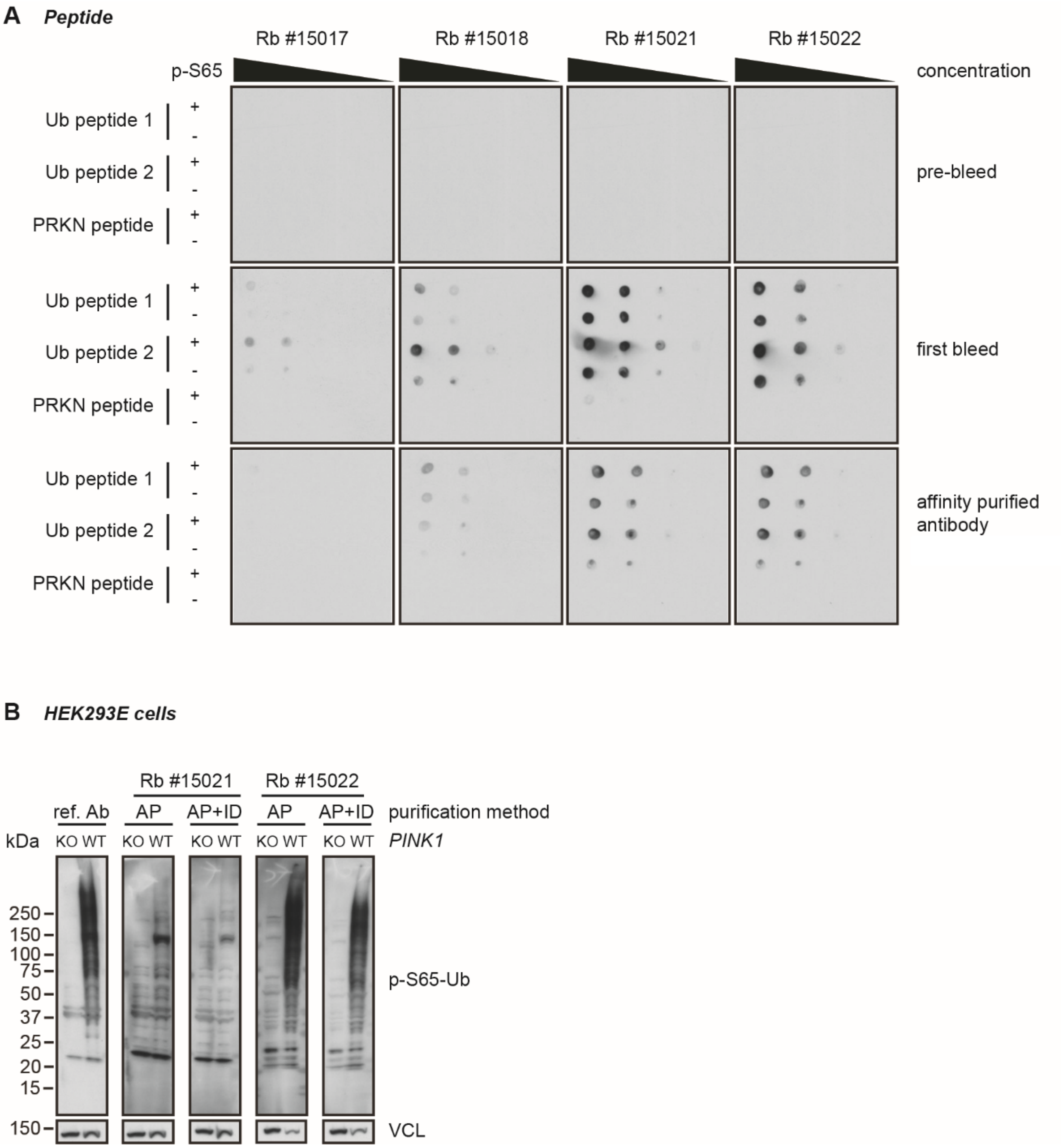


**Supplementary Figure 1. Identification of p-S65-Ub immunopositive bleeds in four rabbits.** (**A**) Dot blot analyses to identify p-S65-Ub positive bleeds from immunized rabbits. Two sets of 13-mer p-S65-Ub peptides containing the p-S65 in its center position that were used for rabbit immunizations as well as negative controls from their non-phosphorylated counterparts and 12-mer non-/phosphorylated PRKN peptides containing the S65 in its center position were spotted on membranes in different concentrations (0.2-25 µM; 5-fold serial dilution). Blots were probed with rabbit sera from before immunization (pre-bleed), after immunization (first bleed), or after affinity-purification. (**B**) WT and *PINK1* KO HEK293E cells were treated for 8 h with 20 µM CCCP and cell lysates were used for western blot analyses. Blots were probed with the reference antibody or bleeds from two rabbits (Rb #15021 and #15022) with either affinity purification (AP) alone or also immuno-depletion (AP+ID). VCL was used as loading control. Ref. Ab - reference antibody, AP - affinity purification, ID - immuno-depletion, KO - knockout, WT - wild-type.

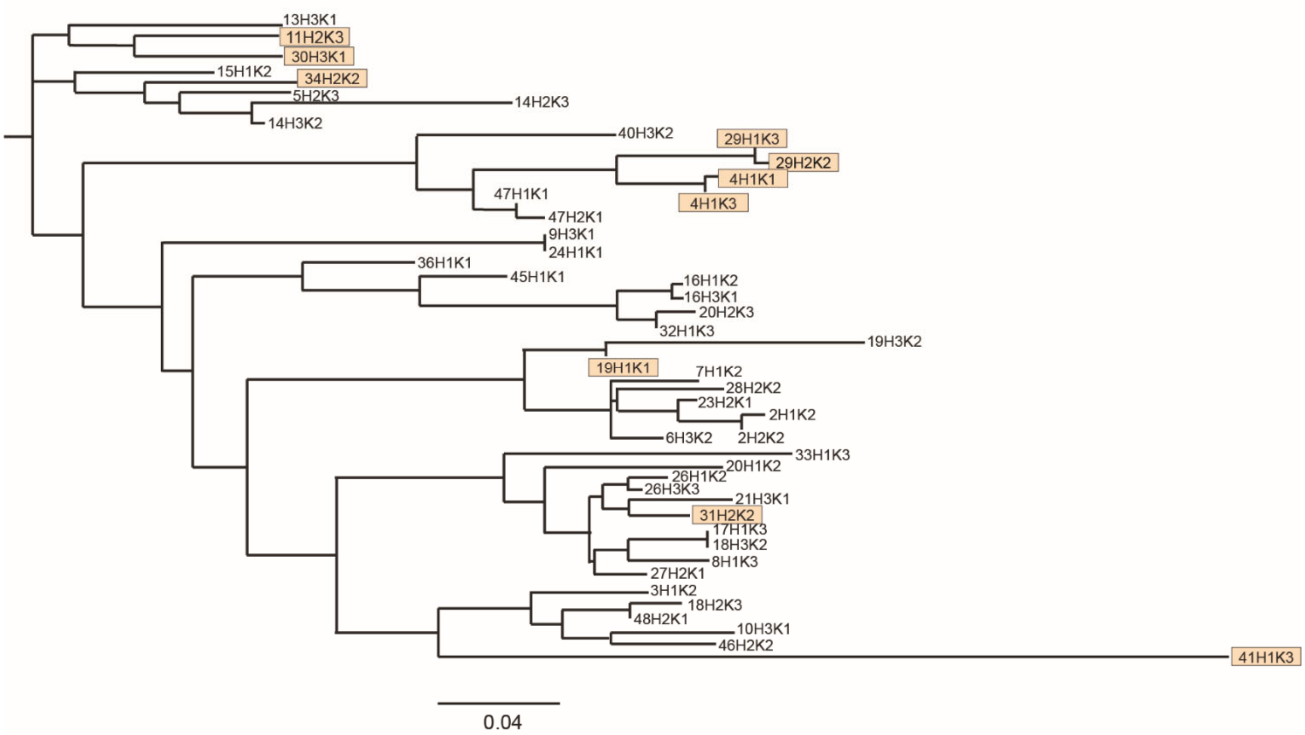


**Supplementary Figure 2. Phylogenetic sequence tree of p-S65-Ub clone variable region.**

A phylogenetic sequence tree was generated based on the p-S65-Ub antibody clone amino acid sequence for combined heavy and kappa variable regions. The top ten promising p-S65-Ub clone supernatants are highlighted in light orange. Scale bar: 0.04 nucleotide substitutions per site.

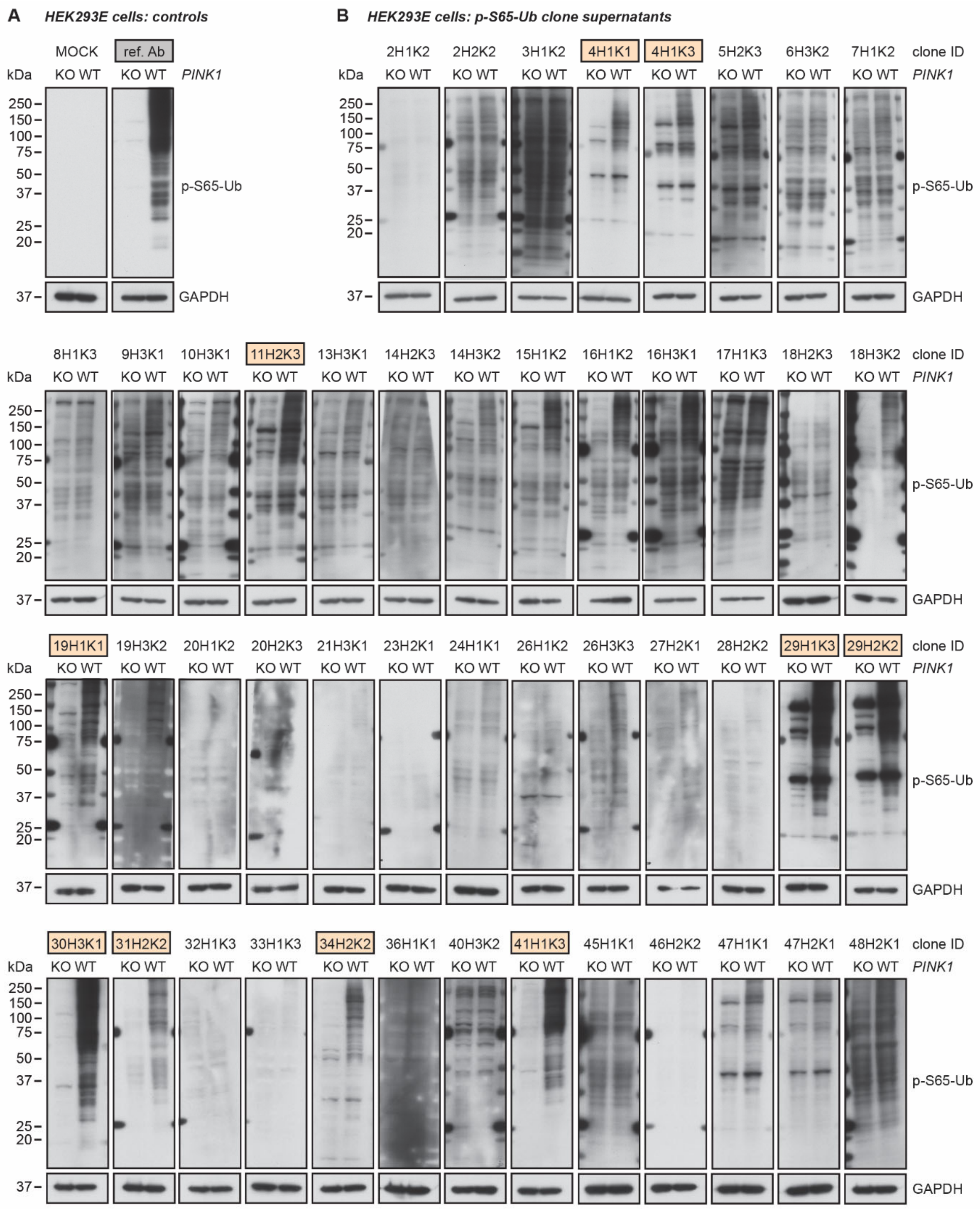


**Supplementary Figure 3. Western blot screening of top recombinant p-S65-Ub clone supernatants.** WT and *PINK1* KO HEK293E cells were treated for 24 h with 20 µM CCCP and cell lysates were used for western blot analyses. (**A**) Representative western blot images from supernatant produced by HEK293 cells transfected with a media control containing all media and transfection reagent except the plasmids (MOCK) vector that served as negative control or from the reference antibody that served as positive control. (**B**) Representative western blot images from 47 p-S65-Ub recombinant antibody supernatants. GAPDH was used as loading control. The top ten promising p-S65-Ub antibody clones are highlighted in light orange. ref. Ab - reference antibody, KO - knockout, WT - wild-type.

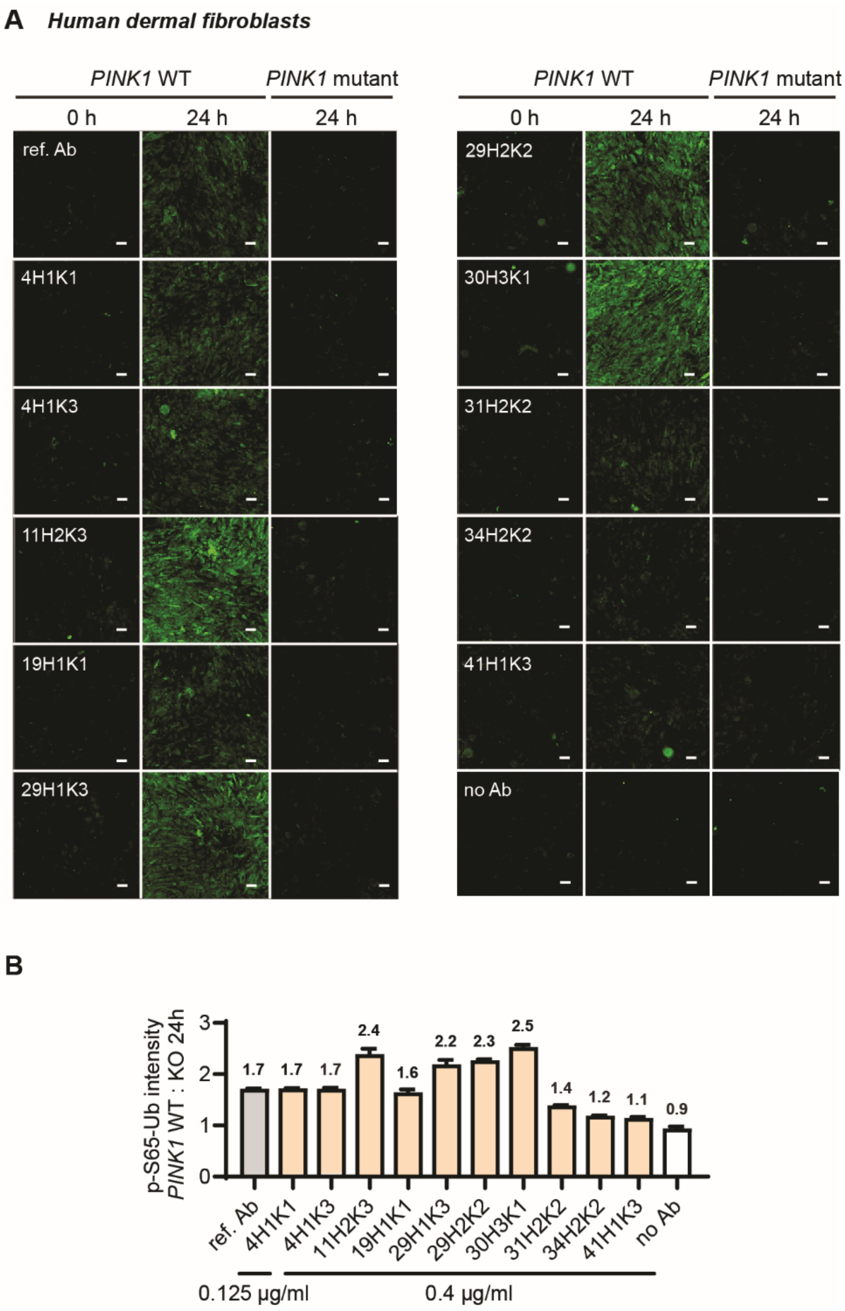


**Supplementary Figure 4. Characterization of the top recombinant p-S65-Ub clone supernatants for immunocytochemistry staining in human dermal fibroblasts.** All ten p-S65-Ub clone supernatants (0.4 µg/ml) and the reference antibody (0.125 µg/ml) were evaluated by immunocytochemistry in fibroblasts treated with 2 µM valinomycin for 0 or 24 hours. (**A**) Representative images of p-S65-Ub immunoreactive signals (green) in human primary skin fibroblasts carrying WT or homozygous *PINK1*$^{Q456X}$ mutation. (**B**) Fluorescence intensities of each clone were quantified by high content imaging and compared relative to the treated *PINK1* mutant fibroblasts. Fold changes are labeled at the top of each bar. N = 2. Samples stained without primary antibody (no Ab) was used a negative control. Scale bar: 50 µm. ref. Ab - reference antibody, WT - wild-type.

**Supplementary Table 1. ELISA screening of B cell supernatants**

| Ranking | B Cell ID | ELISA detection for proteins or peptides | | | | | | Generated recombinant antibody clones |
|---|---|---|---|---|---|---|---|---|
| | | p-S65-Ub protein | Ub protein (non-phospho) | p-S65-Ub peptide | Ub peptide (non-phospho) | p-S65-PRKN peptide | PRKN peptide (non-phospho) | |
| 1 | **19G7[1]** | 2.269 | 0.054 | 1.846 | 0.05 | 0.05 | 0.049 | - |
| 2 | **10F4[1]** | 2.262 | 0.051 | 1.976 | 0.046 | 0.047 | 0.048 | 2H1K2, 2H2K2 |
| 3 | **22A9[1]** | 2.214 | 0.051 | 1.979 | 0.048 | 0.047 | 0.049 | 3H1K2 |
| 4 | **22C1[1]** | 2.192 | 0.056 | 1.821 | 0.048 | 0.048 | 0.048 | 4H1K1, 4H1K3 |
| 5 | **40C7[1]** | 2.114 | 0.047 | 1.794 | 0.05 | 0.05 | 0.051 | 5H2K3 |
| 6 | **4B3[1]** | 2.107 | 0.046 | 1.472 | 0.048 | 0.048 | 0.053 | 6H3K2 |
| 7 | **44B5[1]** | 2.104 | 0.053 | 1.909 | 0.161 | 0.049 | 0.056 | 7H1K2 |
| 8 | **11G6[1]** | 2.085 | 0.051 | 1.931 | 0.05 | 0.052 | 0.049 | 8H1K3 |
| 9 | **14D10[1]** | 2.049 | 0.053 | 1.62 | 0.049 | 0.049 | 0.049 | 9H3K1 |
| 10 | **2E1[1]** | 2.045 | 0.049 | 1.577 | 0.052 | 0.047 | 0.06 | 10H3K1 |
| 11 | **22H5[1]** | 2.016 | 0.05 | 1.92 | 0.051 | 0.047 | 0.055 | 11H2K3 |
| 12 | **44D8[1]** | 2.003 | 0.048 | 2.005 | 0.132 | 0.051 | 0.052 | - |
| 13 | **5A6[1]** | 1.988 | 0.049 | 1.591 | 0.047 | 0.048 | 0.05 | 13H3K1 |
| 14 | **24H10[1]** | 1.986 | 0.051 | 1.51 | 0.052 | 0.05 | 0.051 | 14H2K3, 14H3K2 |
| 15 | **44D11[1]** | 1.975 | 0.05 | 1.866 | 0.165 | 0.05 | 0.052 | 15H1K2 |
| 16 | **15F8[1]** | 1.974 | 0.05 | 1.594 | 0.049 | 0.048 | 0.049 | 16H1K2, 16H3K1 |
| 17 | **5D6[1]** | 1.963 | 0.049 | 1.661 | 0.051 | 0.048 | 0.055 | 17H1K3 |
| 18 | **41E10[1]** | 1.948 | 0.047 | 1.811 | 0.048 | 0.048 | 0.049 | 18H2K3, 18H3K2 |
| 19 | **1F4[1]** | 1.933 | 0.049 | 1.451 | 0.049 | 0.049 | 0.051 | 19H1K1, 19H3K2 |
| 20 | **4E10[1]** | 1.92 | 0.049 | 1.507 | 0.049 | 0.047 | 0.054 | 20H1K2, 20H2K3 |
| 21 | **14C2[1]** | 1.9 | 0.049 | 1.781 | 0.05 | 0.048 | 0.056 | 21H3K1 |
| 22 | **23C11[1]** | 1.875 | 0.05 | 1.539 | 0.049 | 0.049 | 0.049 | - |
| 23 | **44E1[1]** | 1.87 | 0.052 | 1.689 | 0.077 | 0.071 | 0.05 | 23H2K1 |
| 24 | **8B3[1]** | 1.865 | 0.047 | 1.635 | 0.049 | 0.052 | 0.05 | 24H1K1 |
| 25 | **9F8[1]** | 1.854 | 0.048 | 1.4 | 0.046 | 0.048 | 0.049 | - |
| 26 | **24A7[1]** | 1.823 | 0.052 | 1.608 | 0.048 | 0.052 | 0.052 | 26H3K3, 26H3K3 |
| 27 | **1D1[1]** | 1.818 | 0.048 | 1.556 | 0.047 | 0.048 | 0.05 | 27H2K1 |
| 28 | **7D6[1]** | 1.817 | 0.051 | 1.311 | 0.052 | 0.049 | 0.058 | 28H2K2 |
| 29 | **3C11[1]** | 1.803 | 0.051 | 1.584 | 0.048 | 0.049 | 0.05 | 29H1K3, 29H2K2 |
| 30 | **34E2[1]** | 1.788 | 0.049 | 1.819 | 0.054 | 0.053 | 0.051 | 30H3K1 |
| 31 | **15E3[1]** | 1.772 | 0.055 | 1.296 | 0.05 | 0.049 | 0.053 | 31H2K2 |
| 32 | **28D5[1]** | 1.771 | 0.052 | 1.009 | 0.048 | 0.049 | 0.049 | 32H1K3 |
| 33 | **15E9[1]** | 1.771 | 0.049 | 0.728 | 0.053 | 0.048 | 0.056 | 33H1K3 |
| 34 | **27E3[1]** | 1.762 | 0.05 | 1.486 | 0.048 | 0.05 | 0.049 | 34H2K2 |
| 35 | **6H7[1]** | 1.758 | 0.069 | 1.229 | 0.182 | 0.049 | 0.053 | - |
| 36 | **28H[1]** | 1.73 | 0.047 | 1.651 | 0.054 | 0.049 | 0.07 | 36H1K1 |
| 37 | **2G5[2]** | 0.635 | 0.105 | 0.084 | 0.048 | 0.047 | 0.049 | - |
| 38 | **5C10[2]** | 0.443 | 0.046 | 0.149 | 0.049 | 0.048 | 0.05 | - |
| 39 | **37G1[2]** | 0.433 | 0.049 | 0.157 | 0.048 | 0.05 | 0.053 | - |
| 40 | **2B1[2]** | 0.423 | 0.048 | 0.052 | 0.048 | 0.048 | 0.047 | 40H3K2 |
| 41 | **3F8[3]** | 1.754 | 0.047 | 1.367 | 0.067 | 1.831 | 0.062 | 41H1K3 |
| 42 | **17G4[3]** | 1.449 | 0.05 | 0.602 | 0.072 | 1.25 | 0.071 | - |
| 43 | **6B8[3]** | 1.072 | 0.065 | 0.45 | 0.053 | 0.711 | 0.053 | - |
| 44 | **21E11[3]** | 0.715 | 0.052 | 0.221 | 0.05 | 0.335 | 0.053 | - |
| 45 | **30G3[4]** | 0.727 | 0.047 | 0.497 | 0.047 | 0.046 | 0.048 | 45H1K1 |
| 46 | **25B9[4]** | 1.023 | 0.049 | 0.68 | 0.046 | 0.047 | 0.047 | 46H2K2 |
| 47 | **31A7[4]** | 1.019 | 0.049 | 0.856 | 0.048 | 0.049 | 0.051 | 47H1K1, 47H2K1 |
| 48 | **37F1[4]** | 0.95 | 0.048 | 0.903 | 0.048 | 0.051 | 0.051 | 48H2K1 |
| | 44A3 | 2.387 | 0.062 | 2.243 | 0.453 | 0.048 | 0.053 | |
| | 44B6 | 2.322 | 0.055 | 2.091 | 0.476 | 0.058 | 0.06 | |
| | 44B3 | 2.292 | 0.057 | 2.217 | 1.157 | 0.059 | 0.077 | |
| | 44B7 | 2.288 | 0.051 | 2.031 | 0.896 | 0.051 | 0.058 | |
| | 5D2 | 2.207 | 0.051 | 1.955 | 0.49 | 0.049 | 0.052 | |
| | 44C6 | 2.202 | 0.051 | 1.956 | 0.35 | 0.05 | 0.055 | |
| | 44C4 | 2.199 | 0.052 | 1.881 | 0.253 | 0.05 | 0.056 | |
| | 5C6 | 2.169 | 0.568 | 1.912 | 0.566 | 0.049 | 0.049 | |
| | 44D2 | 2.117 | 0.065 | 1.929 | 1.472 | 0.061 | 0.07 | |
| | 44D5 | 2.093 | 0.25 | 2.09 | 1.409 | 0.348 | 0.283 | |
| | 44C5 | 2.09 | 0.055 | 1.637 | 0.215 | 0.048 | 0.058 | |

| Ranking | B Cell ID | ELISA detection for proteins or peptides | | | | | | Generated recombinant antibody clones |
|---|---|---|---|---|---|---|---|---|
| | | p-S65-Ub protein | Ub protein (non-phospho) | p-S65-Ub peptide | Ub peptide (non-phospho) | p-S65-PRKN peptide | PRKN peptide (non-phospho) | |
| | 44B4 | 2.083 | 0.05 | 1.952 | 1.48 | 0.048 | 0.053 | |
| | 10G11 | 2.073 | 0.276 | 1.816 | 0.691 | 0.22 | 0.415 | |
| | 44G7 | 2.06 | 0.063 | 2.079 | 0.843 | 0.051 | 0.055 | |
| | 44F9 | 2.048 | 0.052 | 1.765 | 1.547 | 0.049 | 0.051 | |
| | 44A4 | 2.039 | 0.057 | 2.203 | 2.06 | 0.053 | 0.052 | |
| | 44F6 | 2.03 | 0.052 | 1.849 | 1.551 | 0.049 | 0.055 | |
| | 44A1 | 1.942 | 0.081 | 1.97 | 0.459 | 0.051 | 0.054 | |
| | 31B7 | 1.926 | 0.051 | 1.872 | 0.387 | 0.059 | 0.072 | |
| | 44C8 | 1.925 | 0.049 | 2.083 | 1.998 | 0.05 | 0.055 | |
| | 44F8 | 1.906 | 0.05 | 1.809 | 1.433 | 0.05 | 0.052 | |
| | 44F10 | 1.798 | 0.05 | 1.593 | 0.372 | 0.05 | 0.051 | |
| | 31D10 | 1.739 | 0.048 | 1.146 | 0.048 | 0.048 | 0.048 | |
| | 12B6 | 1.728 | 0.048 | 1.253 | 0.048 | 0.049 | 0.049 | |
| | 34H4 | 1.719 | 0.048 | 1.445 | 0.048 | 0.054 | 0.049 | |
| | 18D5 | 1.664 | 0.049 | 1.387 | 0.051 | 0.048 | 0.053 | |
| | 44E9 | 1.661 | 0.05 | 1.945 | 0.197 | 0.055 | 0.063 | |
| | 3H8 | 1.658 | 0.046 | 1.25 | 0.047 | 0.047 | 0.049 | |
| | 44F4 | 1.645 | 0.053 | 1.854 | 1.031 | 1.401 | 1.023 | |
| | 27A6 | 1.621 | 0.064 | 1.413 | 0.05 | 0.049 | 0.048 | |
| | 22A8 | 1.602 | 0.057 | 0.922 | 0.064 | 0.051 | 0.06 | |
| | 39D5 | 1.571 | 0.048 | 1.135 | 0.048 | 0.049 | 0.049 | |
| | 3D1 | 1.55 | 0.047 | 1.103 | 0.048 | 0.05 | 0.049 | |
| | 11H6 | 1.535 | 0.048 | 0.94 | 0.047 | 0.047 | 0.048 | |
| | 6B6 | 1.529 | 0.048 | 1.676 | 0.102 | 0.049 | 0.052 | |
| | 31C2 | 1.528 | 0.047 | 0.961 | 0.049 | 0.049 | 0.051 | |
| | 44H8 | 1.518 | 0.051 | 0.833 | 0.094 | 0.05 | 0.053 | |
| | 44C7 | 1.508 | 0.056 | 1.922 | 2.093 | 0.059 | 0.057 | |
| | 28C4 | 1.507 | 0.051 | 1.023 | 0.053 | 0.048 | 0.052 | |
| | 44F3 | 1.481 | 0.085 | 1.664 | 1.298 | 0.361 | 0.055 | |
| | 44B9 | 1.469 | 0.053 | 1.445 | 0.97 | 0.051 | 0.056 | |
| | 44C11 | 1.466 | 0.052 | 1.585 | 0.775 | 0.054 | 0.055 | |
| | 1B11 | 1.458 | 0.047 | 0.837 | 0.05 | 0.048 | 0.054 | |
| | 25B11 | 1.448 | 0.05 | 0.912 | 0.048 | 0.049 | 0.049 | |
| | 15G10 | 1.438 | 0.049 | 0.981 | 0.049 | 0.05 | 0.05 | |
| | 21B9 | 1.434 | 0.048 | 1.429 | 1.254 | 0.049 | 0.06 | |
| | 13D4 | 1.428 | 0.051 | 0.815 | 0.061 | 0.051 | 0.057 | |
| | 39E5 | 1.415 | 0.047 | 0.549 | 0.048 | 0.048 | 0.05 | |
| | 44A11 | 1.41 | 0.059 | 1.768 | 1.005 | 0.051 | 0.054 | |
| | 44G10 | 1.407 | 0.05 | 1.175 | 0.056 | 0.048 | 0.049 | |
| | 35D10 | 1.395 | 0.048 | 1.419 | 0.05 | 0.05 | 0.047 | |
| | 37G3 | 1.395 | 0.048 | 0.715 | 0.056 | 0.048 | 0.05 | |
| | 32B2 | 1.393 | 0.046 | 1.448 | 0.046 | 0.049 | 0.047 | |
| | 19G9 | 1.373 | 0.05 | 1.046 | 0.049 | 0.049 | 0.049 | |
| | 30H4 | 1.356 | 0.047 | 1.14 | 0.048 | 0.048 | 0.05 | |
| | 1H3 | 1.347 | 0.048 | 0.763 | 0.05 | 0.049 | 0.048 | |
| | 25D10 | 1.342 | 0.051 | 0.904 | 0.049 | 0.052 | 0.051 | |
| | 29C11 | 1.331 | 0.049 | 1 | 0.049 | 0.048 | 0.049 | |
| | 37D10 | 1.326 | 0.047 | 1.389 | 0.05 | 0.054 | 0.049 | |
| | 28D7 | 1.322 | 0.058 | 0.937 | 0.048 | 0.048 | 0.05 | |
| | 2B11 | 1.322 | 0.048 | 0.675 | 0.047 | 0.047 | 0.054 | |
| | 44C1 | 1.31 | 0.052 | 2.059 | 0.053 | 0.05 | 0.049 | |
| | 24A10 | 1.304 | 0.05 | 0.654 | 0.048 | 0.048 | 0.048 | |
| | 13A8 | 1.302 | 0.049 | 0.652 | 0.05 | 0.051 | 0.055 | |
| | 28F11 | 1.3 | 0.047 | 1.21 | 0.05 | 0.047 | 0.055 | |
| | 11B3 | 1.292 | 0.048 | 0.782 | 0.05 | 0.047 | 0.051 | |
| | 33A5 | 1.268 | 0.049 | 0.764 | 0.05 | 0.051 | 0.05 | |
| | 20G5 | 1.245 | 0.049 | 0.631 | 0.047 | 0.049 | 0.053 | |
| | 35D5 | 1.241 | 0.047 | 1.073 | 0.07 | 0.061 | 0.05 | |
| | 22E9 | 1.233 | 0.052 | 0.758 | 0.048 | 0.047 | 0.047 | |
| | 44A2 | 1.163 | 0.055 | 1.709 | 1.69 | 0.054 | 0.133 | |
| | 44E4 | 1.153 | 0.078 | 2.08 | 1.758 | 0.101 | 0.134 | |
| | 3G1 | 1.146 | 0.047 | 0.681 | 0.134 | 0.048 | 0.053 | |
| | 10F10 | 1.141 | 0.049 | 0.771 | 0.049 | 0.077 | 0.05 | |
| | 11C1 | 1.141 | 0.047 | 0.512 | 0.049 | 0.048 | 0.048 | |
| | 32D1 | 1.109 | 0.047 | 0.954 | 0.051 | 0.047 | 0.05 | |
| | 24F4 | 1.106 | 0.047 | 0.687 | 0.048 | 0.049 | 0.048 | |
| | 2H6 | 1.097 | 0.048 | 0.671 | 0.048 | 0.05 | 0.049 | |
| | 22B6 | 1.096 | 0.051 | 0.433 | 0.048 | 0.05 | 0.057 | |

| Ranking | B Cell ID | ELISA detection for proteins or peptides | | | | | | Generated recombinant antibody clones |
|---|---|---|---|---|---|---|---|---|
| | | p-S65-Ub protein | Ub protein (non-phospho) | p-S65-Ub peptide | Ub peptide (non-phospho) | p-S65-PRKN peptide | PRKN peptide (non-phospho) | |
| | 27C9 | 1.069 | 0.048 | 0.88 | 0.047 | 0.048 | 0.049 | |
| | 25H5 | 1.067 | 0.051 | 0.65 | 0.051 | 0.049 | 0.049 | |
| | 40D5 | 1.054 | 0.062 | 0.352 | 0.049 | 0.049 | 0.075 | |
| | 42H10 | 1.041 | 0.046 | 0.516 | 0.052 | 0.051 | 0.05 | |
| | 8H4 | 1.012 | 0.047 | 0.313 | 0.048 | 0.048 | 0.051 | |
| | 44E2 | 0.929 | 0.061 | 1.819 | 1.53 | 0.05 | 0.066 | |
| | 29G9 | 0.91 | 0.047 | 0.347 | 0.05 | 0.047 | 0.048 | |
| | 44H1 | 0.899 | 0.049 | 1.518 | 1.054 | 0.047 | 0.051 | |
| | 12F1 | 0.877 | 0.049 | 0.372 | 0.046 | 0.049 | 0.052 | |
| | 44B10 | 0.861 | 0.054 | 0.571 | 0.056 | 0.05 | 0.049 | |
| | 38A11 | 0.861 | 0.048 | 0.479 | 0.047 | 0.051 | 0.047 | |
| | 9E6 | 0.858 | 0.771 | 1.027 | 1.033 | 0.663 | 1.155 | |
| | 18D8 | 0.854 | 0.05 | 0.215 | 0.048 | 0.049 | 0.048 | |
| | 44F7 | 0.846 | 0.051 | 0.597 | 0.272 | 0.05 | 0.06 | |
| | 21G2 | 0.837 | 0.054 | 0.259 | 0.047 | 0.051 | 0.053 | |
| | 4B4 | 0.823 | 0.05 | 0.334 | 0.047 | 0.048 | 0.048 | |
| | 44H6 | 0.759 | 0.054 | 1.533 | 0.398 | 0.058 | 0.061 | |
| | 44D6 | 0.751 | 0.053 | 1.351 | 0.784 | 1.1 | 0.432 | |
| | 30A10 | 0.712 | 0.047 | 0.37 | 0.048 | 0.048 | 0.048 | |
| | 28H8 | 0.666 | 0.048 | 0.34 | 0.048 | 0.048 | 0.055 | |
| | 44B8 | 0.655 | 0.052 | 1.553 | 0.376 | 0.048 | 0.058 | |
| | 25H10 | 0.629 | 0.048 | 0.244 | 0.046 | 0.049 | 0.047 | |
| | 34G11 | 0.607 | 0.047 | 0.223 | 0.051 | 0.049 | 0.069 | |
| | 35A7 | 0.595 | 0.046 | 0.202 | 0.049 | 0.047 | 0.047 | |
| | 44G8 | 0.576 | 0.05 | 1.236 | 0.89 | 0.54 | 0.068 | |
| | 44E6 | 0.568 | 0.05 | 0.392 | 0.061 | 0.048 | 0.054 | |
| | 44G9 | 0.523 | 0.049 | 1.757 | 1.494 | 0.049 | 0.051 | |
| | 44E5 | 0.507 | 0.053 | 2.122 | 0.446 | 0.05 | 0.052 | |
| | 28C5 | 0.464 | 0.049 | 0.276 | 0.048 | 0.05 | 0.057 | |
| | 30E6 | 0.444 | 0.046 | 0.628 | 0.401 | 0.05 | 0.05 | |
| | 44G3 | 0.437 | 0.061 | 0.177 | 0.077 | 0.055 | 0.054 | |
| | 44H7 | 0.415 | 0.054 | 1.48 | 0.713 | 0.051 | 0.053 | |
| | 25A6 | 0.414 | 0.051 | 0.168 | 0.05 | 0.048 | 0.05 | |
| | 10G5 | 0.391 | 0.051 | 0.17 | 0.049 | 0.13 | 0.053 | |
| | 44G6 | 0.381 | 0.055 | 1.474 | 0.753 | 0.05 | 0.053 | |
| | 44H9 | 0.37 | 0.054 | 0.187 | 0.051 | 0.214 | 0.05 | |
| | 4A11 | 0.355 | 0.047 | 0.101 | 0.046 | 0.049 | 0.048 | |
| | 39B1 | 0.335 | 0.047 | 0.116 | 0.056 | 0.051 | 0.049 | |
| | 31E4 | 0.325 | 0.047 | 0.098 | 0.048 | 0.048 | 0.047 | |
| | 29D6 | 0.296 | 0.047 | 0.15 | 0.049 | 0.047 | 0.047 | |
| | 44C3 | 0.25 | 0.054 | 1.48 | 0.973 | 0.067 | 0.059 | |
| | 5C8 | 0.242 | 0.046 | 0.096 | 0.048 | 0.049 | 0.048 | |
| | 27E4 | 0.233 | 0.051 | 0.133 | 0.051 | 0.048 | 0.049 | |
| | 44E7 | 0.173 | 0.063 | 1.279 | 1.279 | 0.049 | 0.064 | |
| | 25F2 | 0.165 | 0.051 | 0.092 | 0.051 | 0.049 | 0.048 | |
| | 27A8 | 0.153 | 0.051 | 0.083 | 0.046 | 0.047 | 0.05 | |
| | 44G1 | 0.149 | 0.215 | 1.189 | 0.352 | 0.09 | 0.145 | |
| | 44H5 | 0.147 | 0.056 | 1.647 | 0.915 | 0.049 | 0.058 | |
| | 2F6 | 0.143 | 0.047 | 0.696 | 0.812 | 0.065 | 0.194 | |
| | 32F5 | 0.135 | 0.048 | 0.06 | 0.049 | 0.048 | 0.049 | |
| | 44C2 | 0.124 | 0.05 | 0.787 | 0.162 | 0.05 | 0.053 | |
| | 44H10 | 0.12 | 0.05 | 0.083 | 0.049 | 0.048 | 0.048 | |
| | 44F5 | 0.104 | 0.064 | 1.678 | 1.269 | 0.076 | 0.136 | |
| | 44B2 | 0.097 | 0.062 | 0.749 | 0.491 | 0.049 | 0.055 | |
| | 44D7 | 0.081 | 0.056 | 0.766 | 0.334 | 0.051 | 0.052 | |
| | 44G11 | 0.069 | 0.083 | 0.823 | 0.277 | 0.104 | 0.122 | |
| | 44H2 | 0.061 | 0.048 | 1.194 | 0.42 | 0.047 | 0.05 | |
| | 44A8 | 0.06 | 0.055 | 1.718 | 1.216 | 0.077 | 0.079 | |
| | 44A7 | 0.06 | 0.095 | 0.494 | 0.164 | 0.053 | 0.053 | |
| | 44E3 | 0.059 | 0.05 | 0.425 | 0.05 | 0.049 | 0.05 | |
| | 44E10 | 0.059 | 0.059 | 0.898 | 0.237 | 0.053 | 0.055 | |
| | 44B11 | 0.056 | 0.054 | 1.695 | 1.533 | 0.057 | 0.073 | |
| | 44D1 | 0.054 | 0.05 | 1.109 | 1.09 | 0.05 | 0.051 | |
| | 44H4 | 0.054 | 0.052 | 0.237 | 0.113 | 0.048 | 0.051 | |
| | 44D3 | 0.053 | 0.05 | 1.757 | 1.686 | 0.051 | 0.056 | |
| | 44F11 | 0.052 | 0.051 | 2.096 | 0.544 | 0.052 | 0.058 | |
| | 44D10 | 0.052 | 0.051 | 1.615 | 0.258 | 0.05 | 0.052 | |
| | 44B1 | 0.052 | 0.049 | 1.422 | 1.055 | 0.073 | 0.115 | |

| Ranking | B Cell ID | ELISA detection for proteins or peptides | | | | | | Generated recombinant antibody clones |
|---|---|---|---|---|---|---|---|---|
| | | p-S65-Ub protein | Ub protein (non-phospho) | p-S65-Ub peptide | Ub peptide (non-phospho) | p-S65-PRKN peptide | PRKN peptide (non-phospho) | |
| | 44D4 | 0.052 | 0.052 | 1.09 | 0.967 | 0.05 | 0.056 | |
| | 44F1 | 0.052 | 0.051 | 0.874 | 0.545 | 0.051 | 0.063 | |
| | 44F2 | 0.052 | 0.052 | 0.941 | 0.398 | 0.047 | 0.055 | |
| | 44H3 | 0.052 | 0.049 | 0.196 | 0.092 | 0.048 | 0.113 | |
| | 14E7 | 0.052 | 0.052 | 0.047 | 0.047 | 0.052 | 0.048 | |
| | 44G4 | 0.051 | 0.052 | 1.726 | 0.976 | 0.05 | 0.052 | |
| | 44A6 | 0.051 | 0.053 | 1.227 | 0.373 | 0.056 | 0.065 | |
| | 44H11 | 0.051 | 0.05 | 1.136 | 0.489 | 0.049 | 0.051 | |
| | 44G5 | 0.05 | 0.053 | 1.877 | 0.775 | 0.049 | 0.056 | |
| | 39G9 | 0.05 | 0.051 | 0.048 | 0.046 | 0.048 | 0.048 | |
| | 44E8 | 0.049 | 0.057 | 1.777 | 0.979 | 0.069 | 0.085 | |
| | 44E11 | 0.049 | 0.051 | 1.683 | 1.547 | 0.049 | 0.056 | |
| | 44G2 | 0.049 | 0.072 | 1.195 | 0.121 | 0.049 | 0.05 | |
| | 44D9 | 0.049 | 0.049 | 0.116 | 0.059 | 0.05 | 0.052 | |
| | 44C9 | 0.049 | 0.05 | 0.091 | 0.065 | 0.049 | 0.082 | |
| | 44C10 | 0.048 | 0.05 | 0.272 | 0.049 | 0.048 | 0.049 | |
| | 44A10 | 0.048 | 0.049 | 0.13 | 0.047 | 0.05 | 0.049 | |
| | 44A5 | 0.048 | 0.049 | 0.158 | 0.072 | 0.049 | 0.051 | |
| | 39G11 | 0.048 | 0.063 | 0.047 | 0.048 | 0.049 | 0.058 | |
| | 44A9 | 0.047 | 0.05 | 1.48 | 0.269 | 0.053 | 0.051 | |

Ub - ubiquitin, "-" - clone not available.

[1] B cells that detect both p-S65-Ub protein and peptide.

[2] B cells that only detect p-S65-Ub protein but not peptide.

[3] B cells that detect both p-S65-Ub protein and peptide but additionally also recognize p-S65-PRKN peptide.

[4] B cells that weakly detect p-S65-Ub protein and peptide.
